# Neprosin belongs to a new family of glutamic peptidase based on *in silico* evidence

**DOI:** 10.1101/2022.02.22.481544

**Authors:** Tiew-Yik Ting, Anis Baharin, Ahmad Bazli Ramzi, Chyan-Leong Ng, Hoe-Han Goh

## Abstract

Neprosin was first discovered in the insectivorous tropical pitcher plants of *Nepenthes* species as a novel protease with prolyl endopeptidase (PEP) activity. Neprosin has two uncharacterized domains of neprosin activation peptide and neprosin. A previous study has shown neprosin activity in hydrolyzing proline-rich gliadin, a gluten component that triggers celiac disease. In this study, we performed *in silico* structure-function analysis to investigate the catalytic mechanism of neprosin. Neprosin sequences lack the catalytic triad and motifs of PEP family S9. Protein structures of neprosins from *Nepenthes* × *ventrata* (NvNpr) and *N. rafflesiana* (NrNpr1) were generated by *ab initio* methods and comparatively assessed to obtain high-quality models. Structural alignment of models to experimental structures in the Protein Data Bank (PDB) found a high structural similarity to glutamic peptidases. Further investigations reveal other resemblances to the glutamic peptidases with low optimum pH that activates the enzyme via autoproteolysis for maturation. Two highly conserved glutamic acid residues, which are stable according to the molecular dynamics simulation, can be found at the active site of the substrate cleft. Protein docking demonstrated that mature neprosins bind well with potent antigen αI-gliadin at the putative active site. Taken together, neprosins represent a new glutamic peptidase family, with a putative catalytic dyad of two glutamic acids. This study illustrates a hypothetical enzymatic mechanism of the neprosin family and demonstrates the useful application of an accurate *ab initio* protein structure prediction in the structure-function study of a novel protein family.

## 1. Introduction

Proteases are a class of enzymes that catalyze proteolysis, a process that breaks down peptide bonds of protein into smaller polypeptides or amino acids. Proteases are involved in various biological processes, ranging from specific proteolysis in the living organism to unspecific proteolysis of food proteins. For instance, proteases play important roles in protein localization, modulation of protein-protein interactions, molecular signal transduction, and generation of cell information (Barrett 2001). To date, the MEROPS peptidase database recorded a total of 281 protein families and 66 clans of peptidase under nine catalytic types, namely aspartic (clan A-) cysteine (clan C-), glutamic (clan G-), metallo (clan M-) asparagine (clan N-), mixed (clan P-) serine (clan S-), threonine (clan T-), and unknown catalytic type (clan U-) (Rawlings et al. 2018). Proteases are popular enzymes of interest as they have wide applications in many fields, such as industrial, medical, and biological studies due to their wide substrate specificity and complex functions.

Proteases have been reported to be secreted into the pitcher fluids of *Nepenthes*, which is a genus in the Nepenthaceae tropical carnivorous pitcher plant family with over 170 species that are studied to understand the physiology and mechanisms of botanical carnivory (Clarke et al. 2018). With high-throughput technology, omics-based explorations have been conducted on the *Nepenthes* pitcher tissues and fluids for biomolecular discovery and molecular physiology studies based on transcriptomics, proteomics, and metabolomics approaches (Ravee et al. 2021; Rosli et al. 2021). Proteases in the pitcher fluids have unique characteristics such as higher enzyme stability in a wider range of temperature and pH conditions (Ravee et al. 2018). Therefore, proteases from *Nepenthes* species have great potential for industrial applications. Proteases that have been reported include the aspartic proteinase nepenthesin (Amagase 1972; Amagase et al. 1969) and the neprosin (Lee et al. 2016). Neprosin is reported as a novel peptidase with prolyl endopeptidase activity, which comprises two uncharacterized domains (Lee et al. 2016). The two domains that are previously denoted as the domain of unknown function (DUF) are the neprosin activation peptide (Neprosin_AP) domain (PF14365, previously DUF4409) or neprosin_propep (IPR025521) and the neprosin domain (PF03080, previously DUF239).

Neprosin was first discovered in *N*. × *ventrata* guided by the transcriptome sequences of *N. rafflesiana* (Lee et al. 2016). Neprosins were subsequently profiled in *N. ampullaria* and *N. rafflesiana* via proteomics informed by a transcriptomics study of pitchers and fluids (Wan Zakaria et al. 2016; Zulkapli et al. 2021; Zulkapli et al. 2017). Endogenous neprosins purified from the pitcher fluids of one thousand *N*. × *ventrata* pitchers showed prolyl endopeptidase activity (Rey et al. 2016). This low molecular mass (<40 kDa) neprosin preferentially cleaves at the C-terminal proline residues under acidic (pH 2-3) conditions and can digest protein even at low concentrations without substrate size restriction. The study demonstrated the ability of neprosin and nepenthesin from *N*. × *ventrata* in digesting gliadin, an allergenic gluten component that can trigger celiac disease (Rey et al. 2016). Neprosin has also been explored as a tool for mass spectrometry in bottom-up proteomics studies and histone mapping (Schräder et al. 2017).

Despite many reports on the pitcher fluid protein profiling (Rottloff et al. 2016; Wan Zakaria et al. 2018), only a few proteases among the aspartic proteases, neprosins, cysteine proteases, and serine carboxypeptidases found in *Nepenthes* species have been characterized (Buch et al. 2015; Ravee et al. 2018). The low concentrations of proteases in pitcher fluids hinder large-scale protein purification for functional characterization. Rey et al. (2016) used up to 5 liters of pitcher fluids from 1,000 pitchers fed with flies for six months to obtain sufficient enzymes to study neprosin activity. Neprosin has been categorized into a family with an unknown catalytic type (U74) in the MEROPS protease database (Rawlings et al. 2018). One reason is the lack of neprosin protein structure for inference of catalytic mechanism and biological function with unknown active cleft and catalytic residues. Moreover, there is no experimental evidence for the catalytic mechanism of the neprosin. Hence, this study aims to address this knowledge gap via extensive *in silico* sequence analysis, *ab initio* protein structure modeling, protein docking, and molecular dynamics (MD) simulation. The high-quality protein structures shed light on the catalytic activity of neprosins via the two conserved residues of glutamic acids at the active site, which is akin to the glutamic peptidase family G3.

## 2. Methods

### 2.1. Retrieval of neprosin sequences

*Nepenthes ampullaria* and *N. rafflesiana* samples in this study originated from the population in Taman Negara Endau Rompin, Malaysia. *N. ampullaria and N. rafflesiana* samples were obtained from the UKM *Nepenthes* plot (GPS coordinate: 2°55’11.5” N 101°47’01.4” E). Based on the *N*. × *ventrata* neprosin sequence (NvNpr) reported by Rey et al. (2016), the Basic Local Alignment Search Tool for Protein (BLASTP) analysis was performed to identify homologous neprosin sequences of *N. ampullaria* (NaNpr) and *N. rafflesiana* (NrNpr) based on the recent transcriptome profiling studies (Goh et al. 2020; Zulkapli et al. 2021). NaNpr amino acid sequences were retrieved from the NCBI GenBank database using the accession ID ARA95695.1 and ARA95696.1, whereas NrNpr amino acid sequences were discovered from the transcriptome data of PacBio sequencing (Zulkapli et al. 2017).

### 2.2. In silico analysis of neprosin sequences

SignalP 4.1 (Nielsen 2017) was used to predict signal peptides of neprosin proteins. PfamScan and InterPro were used to identify the protein domains of neprosins. DiANNA 1.1 webserver was used to predict the disulfide bonds in neprosin (Ferrè et al. 2006). EMBOSS Pepstats (Madeira et al. 2019) was used to predict the molecular mass and isoelectric point (pI) of neprosin with default settings. The BLASTP search against NCBI nr database was carried out to find amino acid sequences with high identity. Phylogenetic tree analysis with maximum likelihood method (MLM) was performed for 500 bootstrap replicates using MEGAX with the top 10 hits of sequences from the BLASTP results of the neprosins from *N*. × *ventrata* (NvNpr), *N. rafflesiana* (NrNpr1 and NrNpr2), and *N. ampullaria* (NaNpr1 and NaNpr2). BLASTP analysis was also performed using Dicots Plaza 5.0 against the *Arabidopsis thaliana* genome sequence database (Van bel et al. 2022). The conserved amino acids in neprosins were determined via a multiple sequence alignment (MSA) using neprosins and their BLASTP hits against NCBI nr and Dicots Plaza 5.0 *Arabidopsis thaliana* genome sequence database with Clustal Omega (Madeira et al. 2019) and ConSurf (Ashkenazy et al. 2016). The workflow of the *in silico* analysis is summarized in Fig. S1.

### 2.3. Neprosin protein ab initio structural prediction

The predicted signal peptides were excluded from the structure prediction. The three-dimensional (3D) structures of neprosin protein were predicted using RoseTTAFold (Baek et al. 2021) and AlphaFold2 with MMseqs2 (denoted as AlphaFold2 in the following description) via ColabFold in Google Colaboratory (Jumper et al. 2021; Mirdita et al. 2021). RoseTTAFold is available at the Robetta server (https://robetta.bakerlab.org). The amino acid sequences of neprosin were used as the input and the RoseTTAFold option was selected with default settings before job submission for structure prediction. AlphaFold2 with MMseqs2 was accessible on Google Colab (https://colab.research.google.com/github/sokrypton/ColabFold/blob/main/AlphaFold2.ipynb). Amino acid sequences of neprosins were input as “query_sequence” with default settings. The Protein Data Bank (PDB) files of rank 1 models of neprosins generated by RoseTTAFold and AlphaFold2 can be downloaded from Figshare (https://doi.org/10.6084/m9.figshare.19187252). The structure assessment of neprosin protein models was carried out using SWISS-MODEL (Waterhouse et al. 2018). Default values or parameters were used for all analyses unless specified otherwise. Secondary structures of predicted models were analyzed by FirstGlance in Jmol (http://firstglance.jmol.org) online server.

### 2.4. Protein structure alignment

The PDB files of AlphaFold2 neprosin models with the highest quality were uploaded to the DALI server (Holm 2020) for PDB search of proteins with similar structures. The online mTM-align tool (Dong et al. 2018) was used for the pairwise protein structure alignment to assess protein structural similarity. The superposition of neprosin protein models was carried out using the “Matchmaker” function in ChimeraX 1.2.5 for the visualization of protein models (Pettersen et al. 2021).

### 2.5. Neprosin pocket prediction and protein docking with αI-gliadin

The PDB files of NvNpr and NrNpr1 protein models and their mature counterparts generated using AlphaFold2 were uploaded to the CASTp3.0 webserver (Tian et al. 2018) to predict binding pockets and Richard’s solvent-accessible surface area and volume. The three largest pockets were selected for each model and color-coded based on Richard’s solvent-accessible surface volume. For protein-substrate docking, a crystal structure of the left-handed polyproline II (PPII) helical αI-gliadin (deaminated) was used as the ligand (Kim et al. 2004). The protein model of αI-gliadin (PFPQPELPY) was extracted from the crystal structure of 1S9V in the PDB database. Neprosin protein models were uploaded to the PatchDock webserver (Schneidman-Duhovny et al. 2005) as receptor molecules, whereas αI-gliadin was uploaded as a ligand molecule. PatchDock is based on a geometric-based molecular docking algorithm with a geometric docking score, thus a higher docking score is preferable. Protein docking models with the highest docking score were chosen for visualization and further analysis in ChimeraX.

### 2.6. Molecular dynamics simulation

Molecular dynamics (MD) simulation of the neprosin models was carried out with the CABS-flex 2.0 webserver (Kuriata et al. 2018). The PDB files of neprosin models were analyzed using default settings (C-alpha distance restraints with CABS generated restraints, protein rigidity: 1.0, number of cycles: 50, cycles between trajectory: 50, and temperature at T=1.40, where T=1.0 is close to the temperature of crystal (native state), T=2.0 enables the complete unfolding of unrestrained small protein chains). Three outputs, namely models, contact maps, and fluctuation plots were generated for each neprosin. The fluctuation plot with amino acids position (X-axis) and root mean square fluctuation (RMSF) after global superposition (Y-axis) shows the stability of each amino acid according to movement distance during the MD simulation.

## 3. Results and Discussion

### 3.1. Neprosin homologs of unknown function are widespread in plants

To date, the InterPro database recorded a total of 4,201 protein entries with a domain architecture (O23359) of Neprosin_AP domain followed by the neprosin domain (PF0308), which were predominately found in plants (4,193), apart from the eight bacterial proteins. According to the protein families database Pfam 35.0 (Mistry et al. 2021), the Neprosin_AP domain (PF14365) was present in 2,509 sequences of 129 plant species and two sequences in two bacterial species compared to the neprosin domain (PF03080), which was present in 3,373 protein sequences of 130 plant species, 41 sequences in 28 fungal species, and 50 sequences in 39 bacterial species as of February 2022. There are 56 sequences containing the neprosin domain in *Arabidopsis thaliana*, many of which are putatively annotated as carboxyl-terminal proteinase, tRNA-splicing ligase, NEP-interacting protein with unknown functions (DUF239), or uncharacterized protein. This suggests the importance of the neprosin family in diverse taxa with yet undiscovered biological functions.

### 3.2. Sequence analysis of neprosins from Nepenthes species

SignalP 4.1 predicted the presence of signal peptides in all neprosins of *Nepenthes* species (Fig. 1). PfamScan and InterPro detected the presence of both the neprosin activation domain (PF14365/IPR025521) and the neprosin domain (PF03080/IPR004314) in neprosins. All neprosins have three disulfide bonds, except NrNpr2 with two disulfide bonds. The reported NvNpr with PEP activity shared the highest sequence identity (96%) with NrNpr1 and the lowest identity (42%) with the longer sequence of NaNpr1.

**Figure 1.**
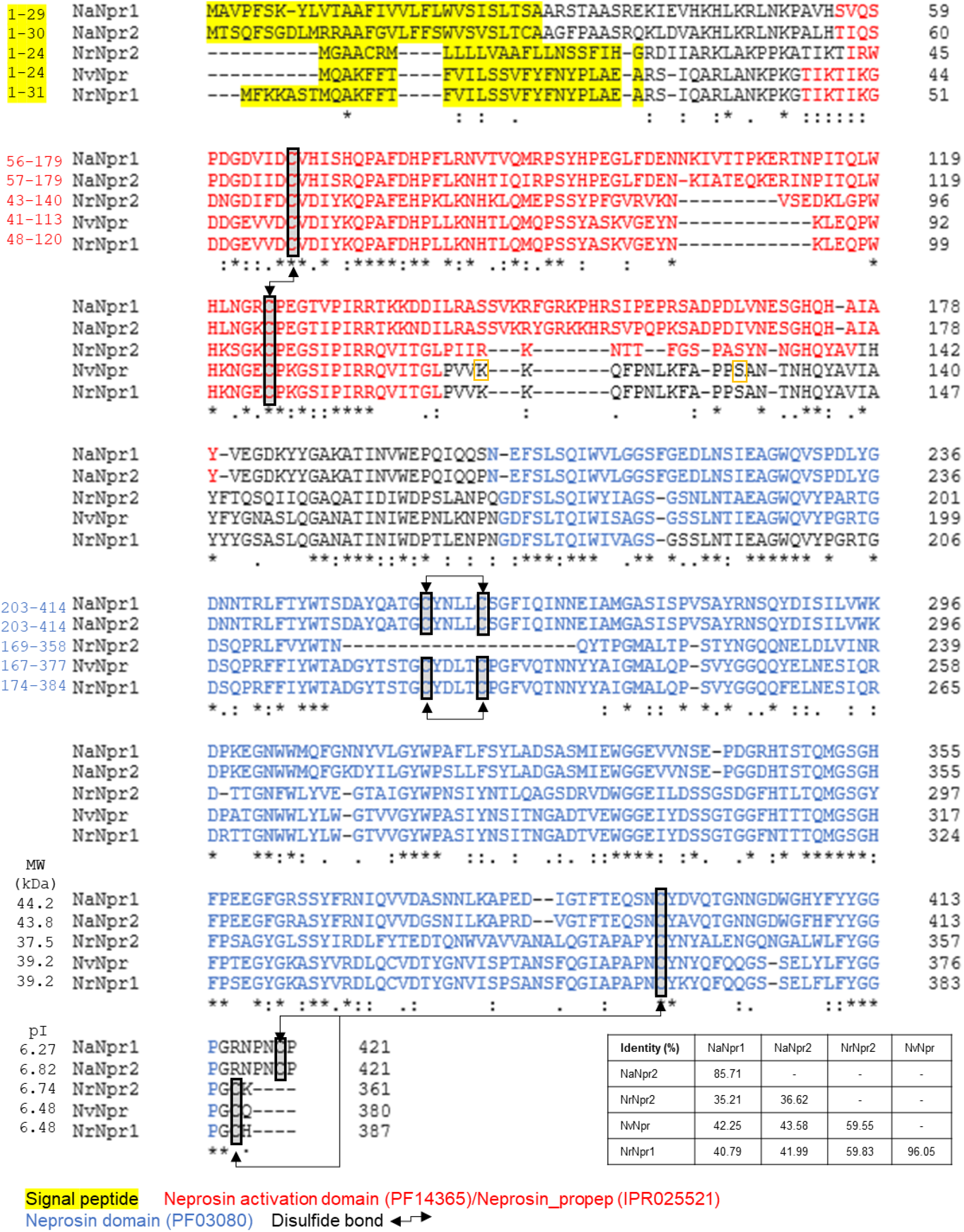
Multiple sequence alignment of neprosins identified from *N*. × *ventrata* (NvNpr, MER0351045), *N. ampullaria* (NaNpr1 and NaNpr2, GenBank: ARA959695.1 and GenBank: ARA9596.1), and *N. rafflesiana* (NrNpr1 and NrNpr2) showing the signal peptide, functional domains, and disulfide bond pairing of cysteine [C] residues. The percentage identity matrix of full-length neprosin amino acid sequences is depicted. The numbers on the left corresponds to the positions of residues for each domain. Theoretical molecular weight (MW) and isoelectric point (pI) are shown for sequences excluding the signal peptide. Two putative start sites for the mature enzyme of NvNpr (Rey et al. 2016) are indicated by orange boxes. The pI for mature NvNpr (m-NvNpr) from the first start site (28.95 kDa) was predicted as 4.63 compared to m-NvNpr from the second start site (27.55 kDa) of 4.30.

A phylogenetic tree was constructed using MEGAX based on the top 10 BLASTP hits of five neprosins from *Nepenthes* species (Fig. S2). Most of the hits were hypothetical, predicted, or uncharacterized proteins. The BLASTP hits of NvNpr, NrNpr1, and NrNpr2 shared a C-terminal peptidase from *Nepenthes alata* (BAW35437.1). In addition to BAW35437.1, NrNpr2 found hits to another *N. alata* C-terminal peptidase (BAW35438.1). NaNpr1 and NaNpr2 BLASTP results contain carboxyl-terminal peptidase, putative metal tolerance protein 4-like, 3-methyl-2-oxobutanoate hydroxymethyltransferase, and tRNA-splicing ligase. According to the phylogenetic tree, neprosins from *N. rafflesiana, N*. × *ventrata*, and C-terminal peptidase from *N. alata* shared a closer most recent common ancestor (MRCA) compared to *N. ampullaria*. Hence, neprosins from *N. ampullaria* are more distant in the evolutionary relationship than other *Nepenthes* species.

A BLASTP search against the *Arabidopsis thaliana* amino acid sequences using Dicots Plaza 5.0 found a carboxyl-terminal peptidase (DUF239) (AT3G48230) and a putative NEP-interacting protein (AT5G19170) to be the closest to NvNpr, NrNpr1, and NrNpr2 (Fig. S3). Meanwhile, NaNpr1 and NaNpr2 were closer to two tRNA-splicing ligases (DUF239) (AT5G56530 and AT1G55360) and a putative carboxyl-terminal peptidase (AT3G13510). Based on the phylogenetic trees generated using BLASTP and *Arabidopsis* hits, closely related amino acid sequences of neprosin were selected for a multiple sequence alignment (MSA) using Clustal Omega (Fig. S4). The MSA was used for the ConSurf analysis to determine functionally conserved amino acids in the neprosins (Fig. 2). Based on the ConSurf results, the catalytic triad Ser-Asp-His commonly found in prolyl endopeptidases (MEROPS S9A family) was not detected. Moreover, the motif of S9A (GGSXGGLL, X normally Asn or Ala), S9B (GWSYGGY), S9C (GGSYGG), and S9D (GGHSYGAFMT) were also not found in the neprosin sequences. Therefore, we deduce that neprosin does not belong to the family S9 of prolyl endopeptidase. Due to sequences not related to any known peptidase, neprosin has been classified as family U74 (unknown catalytic type) in MEROPS.

**Figure 2.**
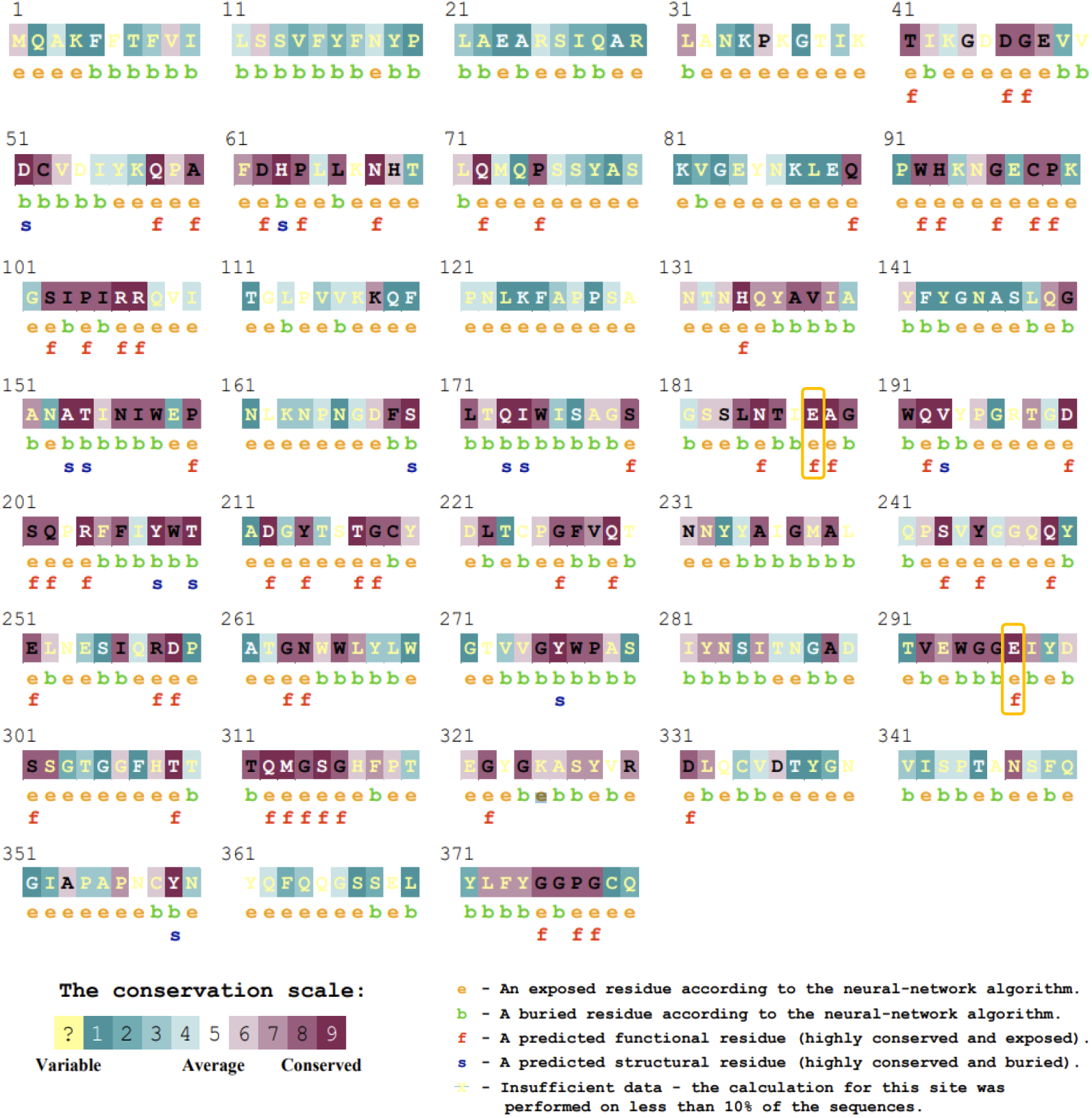
Conservation and functionality of amino acids in the multiple sequence alignment of neprosin family with NvNpr as a query in ConSurf analysis. The orange boxes highlight the highly conserved glutamic acid (E) residues.

### 3.2. Experimental quality neprosin models

For structure prediction to unveil the catalytic mechanism of neprosin, NvNpr was chosen as a reference due to its established prolyl endopeptidase (PEP) activity (Rey et al. 2016). Another homolog from *N. rafflesiana* (NrNpr1) sharing 96% sequence identity with NvNpr (Fig. 1) was included for comparative analysis. Due to the lack of a crystal structure in PDB with significant identity for homology modeling, we applied the two latest *ab initio* methods, namely RoseTTAFold and AlphaFold2, to generate 3D protein structures without structural templates. The first-rank models with the highest confidence level were chosen from the five models generated for each method (Fig. 3). All neprosin models generated were globular proteins with secondary structures mainly comprising coils (38-44%), followed by β-strands (32-34%) and helices (9-12%) (Table S1). The two antiparallel six- and seven-stranded β-sheets form an overall β-sandwich structure like that of glucanases. Notably, independently predicted high-quality neprosin models from both RoseTTAFold and AlphaFold2 showed very high structural similarities, especially in the β-sandwich structure (Fig. 3). RoseTTAFold and AlphaFold2 models of NvNpr can be structurally aligned for 239 amino acids with a mTM-align pairwise TM-score as high as 0.919 and RMSD of 1.920 Å; whereas for NrNpr1 alignment of 322 amino acids, the pairwise TM-score was 0.899 with 2.435 Å RMSD.

**Figure 3.**
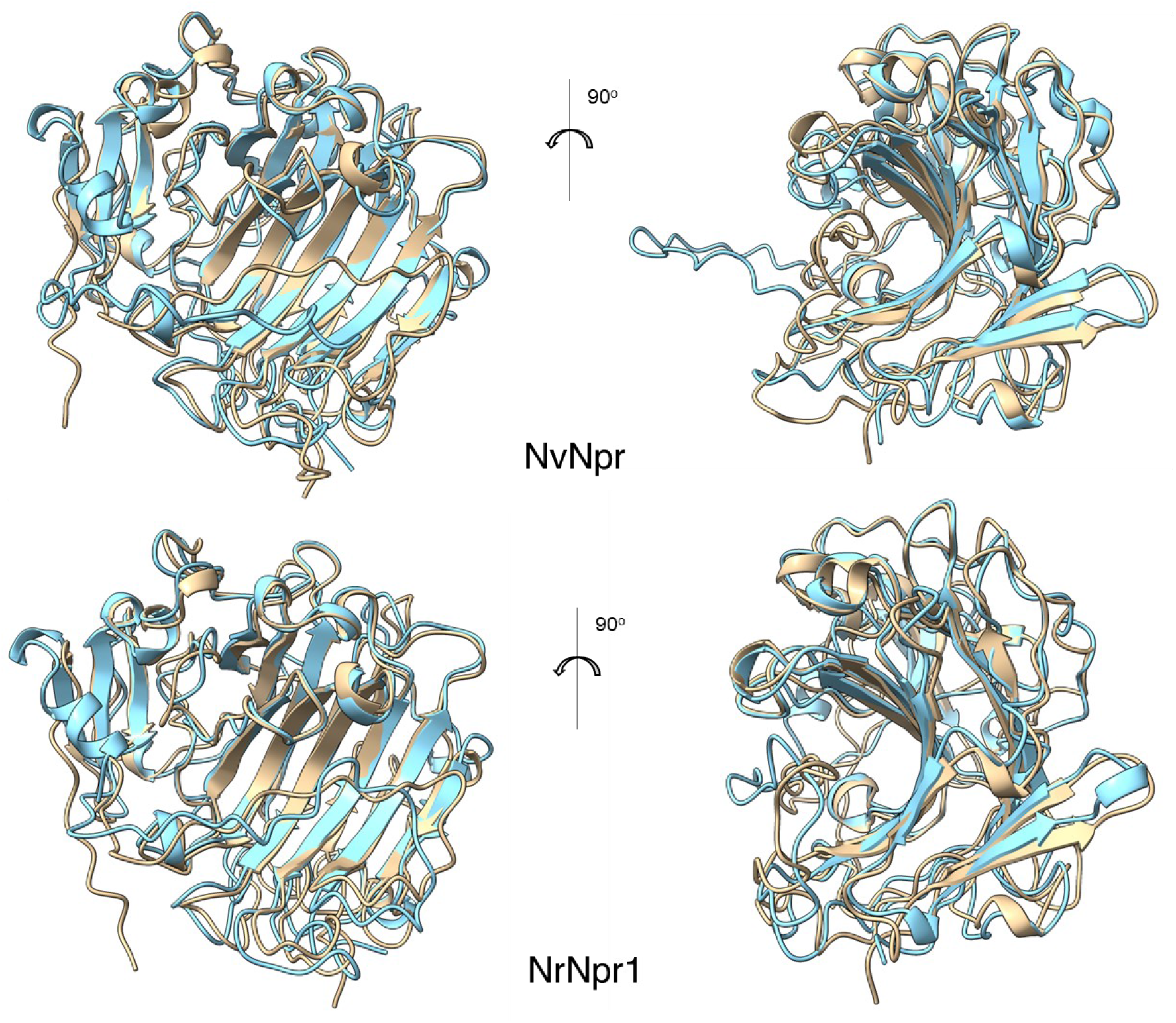
The first-rank models with the best quality generated by AlphaFold2 with MMseqs2 (bronze) and RoseTTAFold (cyan) for NvNpr (top) and NrNpr1 (bottom) superimposed using the ‘Matchmaker’ function in ChimeraX.

Based on the SWISS-MODEL protein model quality assessment (Table 1), Ramachandran favored in neprosin models generated by RoseTTAFold and AlphaFold2 were around 92-96%. Ramachandran favored refers to the fraction of residues in favored regions of the Ramachandran plot, which is ideally >98%. AlphaFold2 models were of higher quality based on the lower clashscores and MolProbity (a composite score of normalized all-atom clashscore, Ramachandran favored, and rotamer outlier). The clashscore is the number of serious steric overlaps (>0.4 Å) per 1,000 atoms and should be ideally close to 0 (Chen et al. 2010). Meanwhile, the QMEANDisCo Global scores of AlphaFold2 models were also higher. Overall, AlphaFold2 outperformed RoseTTAFold in generating near experimental quality neprosin models for further analysis.

**Table 1.** Quality assessment of the first-rank protein models generated by RoseTTAFold and AlphaFold2. Ramachandran favored (%), clashscore, MolProbity score, and QMEANDisCo Global of the models were obtained from structure assessment of SWISS-MODEL. Confidence is assigned to RoseTTAFold models by the Robetta server while pLDDT is provided as an output to models generated by AlphaFold2 with MMseqs2 via ColabFold.

| Method | Model | Ramachandran<br>favored (%) | Clashscore* | MolProbity<br>score* | QMEANDisCo<br>Global | Confidence | pLDDT |
| --- | --- | --- | --- | --- | --- | --- | --- |
| RoseTTAFold | NvNpr | 95.60 | 172.97 | 3.00 | 0.61±0.05 | 0.71 |  |
|  | NrNpr1 | 92.66 | 185.39 | 3.19 | 0.81±0.05 | 0.70 |  |
| AlphaFold2 | NvNpr | 92.66 | 27.68 | 2.39 | 0.75±0.05 |  | 91.2 |
|  | NrNpr1 | 94.07 | 25.82 | 2.30 | 0.81±0.05 |  | 91.6 |

\*Lower score indicates a higher quality

AlphaFold2 models were of higher confidence based on a per-residue measure of local confidence with the overall predicted local distance difference test lDDT-Cα (pLDDT) scores of over 90. A reasonable model has a pLDDT score of 60 or greater, while scores above 80 are great models. Meanwhile, confidence scores of RoseTTAFold models ranged between 0.70 and 0.71. Notably, most of the regions in neprosin models generated using AlphaFold2 showed pLDDT >90, which is expected to be highly accurate and suitable for applications such as characterizing binding sites (Jumper et al. 2021). There were several positions in the models where pLDDT <80 (Fig. 4). At the amino acid position around 100^th^, there was a drop of pLDDT below 40 in both NvNpr and NrNpr1 models. Intriguingly, this position (without signal peptide) matched the second proposed start site of mature NvNpr (Fig. 1). The 90^th^ to 110^th^ amino acids of both NvNpr and NrNpr1 models correspond to the region of amino acids between Neprosin_AP and neprosin domains showed pLDDT <50, suggesting that this region could be unstructured (Fig. 4). Regions with pLDDT <50 often have a ribbon-like appearance and are disordered, either unstructured in physiological conditions or only structured as a part of a complex (Jumper et al. 2021). The disordered region of a protein, also known as the intrinsically disordered protein region (IDPR) provides conformational flexibility and structural dynamics that allow its protein to perform the unorthodox activity that is impossible in ordered proteins (Uversky 2019). The IDPR between the Neprosin_AP and neprosin domains could act as a flexible linker that connects the two structured domains.

**Figure 4.**
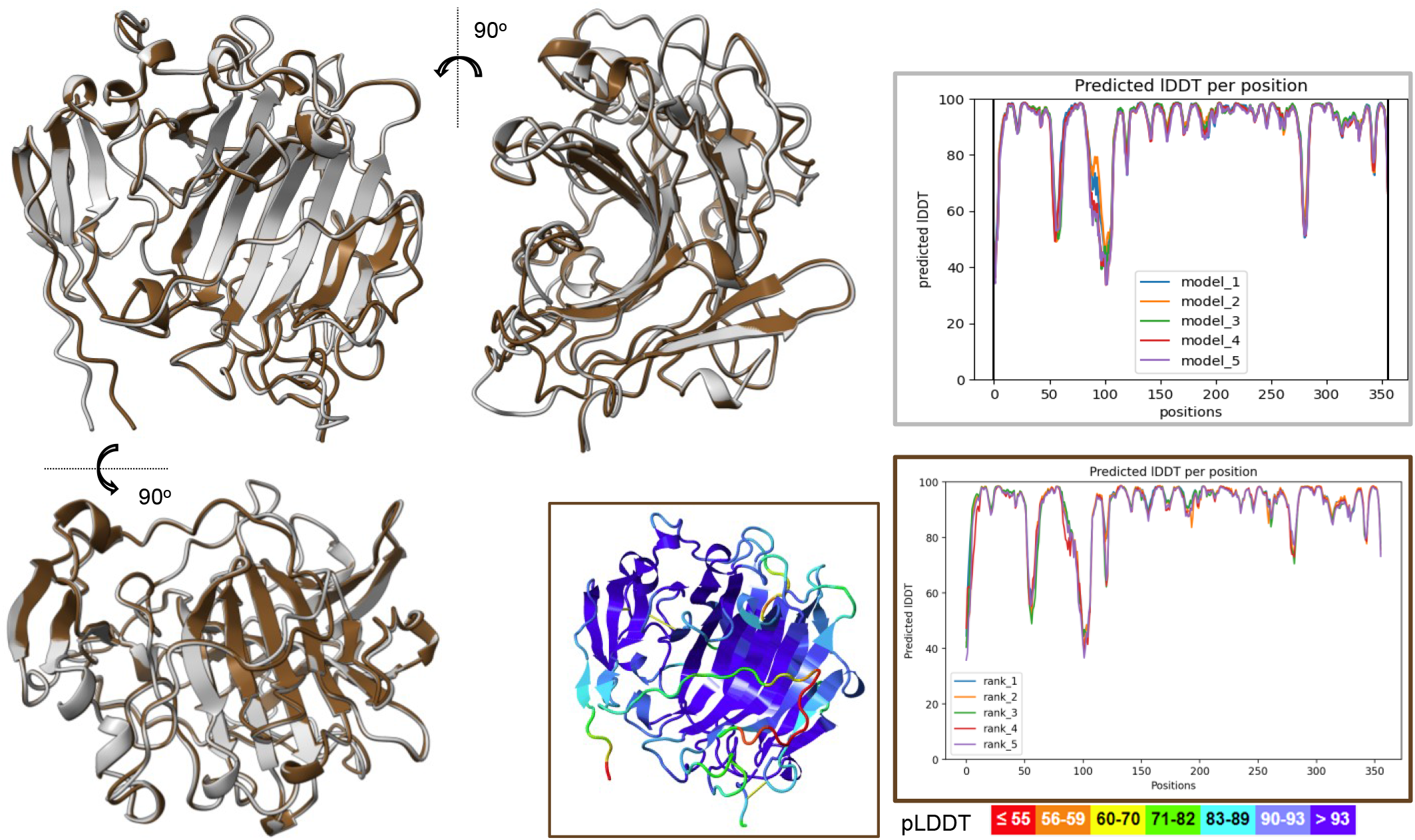
The first-rank AlphaFold2 models of NvNpr (light grey) and NrNpr1 (dark brown) superimposed using ChimeraX ‘Matchmaker’ function with respective pLDDT plots. The inset with dark brown border shows the NrNpr1 model visualized using Jmol FirstGlance and colored according to the pLDDT score.

### 3.3. Protein structural alignment revealed neprosin as a glutamic peptidase

Neprosin protein models from AlphaFold2 were subjected to a heuristic PDB search using the DALI protein structure comparison server. Both NvNpr and NrNpr1 searches (Table 2) found hits to the crystal structure of scytalidoglutamic peptidase or scytalidopepsin B (SGP, PDB ID: 2ifw), which is the founding member of the peptidase family G1, so far found only in fungi. Apart from SGP, DALI search also found chain-B of aspergillopepsin II or aspergilloglutamic peptidase (AGP, PDB ID: 1y43-b) (Table 2).

**Table 2.** Results of DALI search hits in the PDB database. The Z-score, RMSD (Å), sequence identity, and average pairwise length, L_ali_ are assigned to each DALI search hit.

| Attribute | NvNpr |  | NrNpr1 |  |
| --- | --- | --- | --- | --- |
| Hit | 2ifw | 1y43-b | 2ifw | 1y43-b |
| Z-score | 13.6 | 8.9 | 13.5 | 8.9 |
| RMSD (Å) | 2.7 | 3.2 | 2.7 | 3.3 |
| Sequence identity | 0.11 | 0.05 | 0.12 | 0.06 |
| Average pairwise length, $L_{ali}$ | 175 | 149 | 173 | 152 |

SGP and AGP are categorized as members of the glutamic peptidase family G1 in the MEROPS database. Root-mean-square-deviation (RMSD) is a common measurement used for the evaluation of structural similarity but is not used in the DALI scoring. DALI Z-score is a length-dependent rescaling of the collective score of distance matrix alignment and scoring function (Holm 2020). The higher Z-score shows a higher structural similarity between the query and the PDB hits but cannot be used for interpreting homology, as sequence and functional conservation are needed to infer the evolutionary relationship. Holm et al. (2008) define Z-scores above 2 as ‘significant similarities’ and correspond to similar folds. A ‘strong match’ requires either sequence identity > 20% or a Z-score cutoff above (*n*/10)-4 (where *n* is the number of residues in the query structure). The sequence similarities between 2ifw and 1y43-b with neprosins were below 20%, while the Z-score cutoffs for both NvNpr and NrNpr1 were 31.6 [(356/10)-4]. Hence, the hits to glutamic peptidases fall under ‘significant similarities’.

NvNpr and NrNpr1 protein models superimposed with DALI hits of SGP (MEROPS ID: G01.001) and AGP (MEROPS ID: G01:002) crystal structures (Fig. S5) showed very high structural similarity with reported active cleft and catalytic dyad (Fujinaga et al. 2004; Pillai et al. 2007; Sasaki et al. 2004). Interestingly, the neprosin domain structure is very similar to the glucanase-like β-sandwich formed by the two seven-stranded antiparallel β-sheets as described in the eqolisin family or glutamic peptidases (G1). The structure of the neprosin domain resembles the active clefts of glutamic peptidases. This observation supports that the neprosin domain is the catalytic domain with peptidase activity. The glutamic acid residues of the catalytic dyad were found to overlap at the corresponding sites in the active cleft (Fig. S5). Fujinaga et al. (2004) resolved the tertiary structure of SGP and proposed a hydrolytic mechanism, whereby water is bound to Glu136 as the primary catalytic residue while Gln53 acts as a nucleophile after it is activated into hydroxide ion by Glu136 carboxylation. The side-chain amide of Gln53 provides electrophilic assistance for the formation of tetrahedral intermediate and oxyanion stabilization. Site-directed mutagenesis conducted on aspergilloglutamic peptidase (AGP) concurs with the proposed catalytic residues (Yabuki et al. 2004). The mutation of Gln133 (corresponding to Gln53 of SGP) resulted in a complete loss of function of enzymatic activity without changing the conformation of AGP. Furthermore, a Q133E/E219Q double mutant of AGP (Glu219 corresponds to Glu136 of SGP) did not mature via autoproteolysis upon incubation and thus showed no enzymatic activity (Yabuki et al. 2004).

Later, Sasaki et al. (2005) proposed that catalytic glutamic acid of AGP acts as a general acid in the first phase of catalysis by donating H^+^ ion to the carbonyl oxygen of the scissile peptide bond of a substrate. A water molecule donates an OH^-^ group to the carbonyl carbon to form a tetrahedral intermediate. The transition state of the substrate is stabilized by hydrogen bonding with the two catalytic residues. In the next phase, the protonated glutamic acid donates the H^+^ ion to the amide nitrogen atom of the scissile peptide bond, which causes the breakdown of the tetrahedral intermediate and cleavage of the peptide bond.

Both proposed catalytic mechanisms of SGP and AGP suggest the glutamic acid of the catalytic dyad as the general acid that donates the H^+^ ion for the protonation of leaving-group nitrogen, which is an essential step for the hydrolysis of an amide (Fujinaga et al. 2004; Sasaki et al. 2005). Moreover, the solvent kinetic isotope effects and proton inventory studies by Kondo et al. (2010) also support that the catalytic mechanism of SGP requires nucleophilic attack of water molecule activated by a catalytic glutamic residue (Glu136). Meanwhile, the amino acid residue in the neprosin models that is superposed to the catalytic glutamine residue (Gln53 of SGP or Gln24 of AGP) is a glutamic acid (Glu164). Therefore, Glu164 and Glu273 (numbering in NvNpr sequence without signal peptide) were hypothesized as the catalytic dyad responsible for prolyl endopeptidase-like catalytic activity.

The two glutamic acid residues of the hypothetical catalytic dyad were 100% conserved in the neprosin family according to the ConSurf analysis (Fig. 2). The glutamic acid residues (Glu188 and Glu297 for the full-length NvNpr sequences; Glu164 and Glu273 in the NvNpr sequences without the signal peptide) were conserved in all nine sequences. A high conversation of these glutamic acids suggests their important roles in maintaining the structure or function of the neprosin domain. According to InterPro, these two glutamic acid residues were highly conserved in the neprosin domain (PF03080) (Fig. S6). The glutamic acid residues Glu164 and Glu273 were conserved in 211 and 253 of 427 representative proteome 15 (RP15) sequences of the neprosin domain, respectively. Together, the putative catalytic dyad (Glu164 and Glu273) was conserved in 184 (43%) RP15 sequences.

### 3.4. Neprosin belongs to a new family of glutamic peptidase

Based on all the characteristics described above, we found a glutamic peptidase with a catalytic dyad comprising two glutamic acids and post-proline cleaving activity from the MEROPS peptidase family G3. The sole member of the family, strawberry mottle virus (SMoV) glutamic peptidase (MER1365461) has a catalytic dyad of Glu1192 and Glu1274 as proven by a mutagenesis study (Mann et al. 2019). Furthermore, this peptidase appears to have a preference to cleave after proline residue without the limitation of substrate size like that reported in neprosin by Schräder et al. (2017). For instance, SMoV cleaves after proline in a big polyprotein (1,691 amino acids) to form products of peptide-Pro1101+Ala-peptide and peptide-Pro1444+Lys-peptide (Mann et al. 2019). The SMoV peptidase unit ranges from amino acid 1,102^nd^ to 1,335^th^ (234 amino acids), which was modeled using AlphaFold2. The superimposition of this peptidase unit with NvNpr shows the catalytic dyad of this family G3 glutamic peptidase (Glu 91 and Glu 173, numbering in peptidase unit) superposed the hypothetical catalytic dyad of Glu164 and Glu273 in NvNpr and NrNpr1 (Fig. S7). Despite a high structural similarity between neprosins and the SMoV peptidase unit, there is no evidence of sequence homology between them. Therefore, neprosins represent a new family of glutamic peptidase with a similar catalytic mechanism to that of family G3.

### 3.5. Neprosin maturation via autoproteolysis provides substrate access to the active site

Mass spectrometry (MS) analysis by Rey et al. (2016) found that the native NvNpr from the pitcher fluids contains mainly the neprosin domain (PF03080) with an estimated molecular mass of 28.9 kDa. This suggests that neprosin maturation requires proteolysis, which is supported by the observation that recombinant NvNpr required one-week incubation in pH 2.5 buffer for autolytic activation (Schräder et al. 2017). Similarly, mature SGP (260 residues) and AGP (282 residues) have only 206 and 212 amino acids, respectively. The autoproteolytic maturation of AGP has been described by Inoue et al. (1991). The AGP proenzyme comprises 264 amino acids after N-terminal signal peptide (18 residues) removal. The pre-pro sequence (41 residues) for the inhibition and thermal stabilization of the zymogen will be removed in acidic conditions (Inoue et al. 1991; Kubota et al. 2005). Lastly, the 11-residue intervening peptide is removed via autoproteolysis to form the mature AGP (212 residues) with a light chain (39 residues) and a heavy chain (173 residues) that are bound non-covalently.

Another known similarity between neprosin and glutamic peptidases is their optimal working pH. A circular dichroism (CD) spectrum of AGP zymogen dialyzed against buffer at pH 5.25 was identical to that of the full-length recombinant zymogen, while the dialysis of AGP zymogen against pH 3.5 buffer yielded a spectrum identical to that of mature AGP that could digest hemoglobin under acidic (pH 2.0) conditions (Huang et al. 2007). Similarly, SGP maintained its structure and enzymatic activity at pH 2 to 7 but denatured at pH > 8 (Kondo et al. 2010). Meanwhile, NvNpr showed maximum activity at pH 2.5 and active up to pH 5 with near-zero activity at pH 8 (Rey et al. 2016).

Sasaki et al. (2012) proposed a mechanism of autoproteolysis according to the crystal structure of a dimeric AGP. The C-terminal of the light chain of one AGP is found in the active cleft of another AGP like a substrate. This autoproteolytic mechanism could be possible in neprosin as the full-length NvNpr (380 residues) is 42 kDa, about ∼12 kDa heavier than native and active recombinant neprosins with two proposed start sites (Fig. 1). The second start site after a proline residue is more probable for the mature neprosin because no peptide fragment before the “cleaving after P” site was observed in the MS analysis of neprosin by Rey et al. (2016) due to its post-proline cleaving activity (Schräder et al. (2017). The predicted molecular mass for the mature recombinant neprosin (252 amino acids) is 27.6 kDa, which is lighter than the native neprosin with glycosylation.

To investigate neprosin maturation, the start site of mature NrNpr1 was deduced from the sequence alignment with the proposed start sites of mature NvNpr (Fig. 1), which is located within the IDPR (Fig. 4). The flexible IDPR could play a role in neprosin activation via proteolysis by allowing protease binding or conformational changes of neprosins. In AGP, autoproteolysis via the binding of one proenzyme to the active cleft of another resulted in enzyme activation (Sasaki et al. 2012). Such autoproteolysis in neprosin proenzyme could be possible since the IDPR is positioned near the active cleft with high structural similarity to AGP (Fig. 4 & S5). This will need to be experimentally validated by a crystallography study of neprosin.

The predicted mature protein models of NvNpr (m-NvNpr) and NrNpr1 (m-NrNpr1) after the removal of the neprosin activation domain were generated by AlphaFold2 with very high pLDDT scores of 94.4 and 95.1, respectively. Notably, the mature neprosin models showed very high structural similarities to the crystal structures of AGP-chain B and SGP, as well as the AlphaFold2 model of the SMoV peptidase unit (Fig. S8). Based on the mTM-align pairwise structure alignment, the TM-scores between the mature neprosin models (m-NvNpr and m-NrNpr1) and the glutamic peptidases (AGP-chain B, SGP, and SMoV) were above 0.6 despite that the percentage identity was less than 13%. At a TM-score cut-off of 0.5, the P-value is 5.5 × 10^−7^ (1.8 million protein pairs to be achieved by random chance) according to Xu et al. (2010). This means that the structural similarities between glutamic peptidases and mature neprosin were highly significant.

The CASTp 3.0 web server was used to predict empty concavities (pockets) to which solvent can gain access for different neprosin protein models (Fig. 5). In both NvNpr and NrNpr1, the largest pockets were observed in the putative active clefts only in the mature neprosin models. Hence, we hypothesize that the removal of the neprosin activation domain exposes the putative cleft for the activation of neprosin, allowing access for substrate catalysis. A non-physiological substrate of neprosin is αI-gliadin, which is resistant to gastrointestinal digestion while transglutaminase (TG2) deamination confers enhanced immunogenicity in triggering celiac disease by binding to HLA-DQ2 in the intestine (Kim et al. 2004). To investigate whether neprosins can bind with αI-gliadin, the docking of neprosin models with αI-gliadin was conducted using PatchDock. Protein docking with αI-gliadin found no solution in the putative active cleft of neprosin with the presence of the neprosin activation domain. The first docking solutions with high docking scores of 7,752 and 7,956 were found for both mature neprosin models of NvNpr and NrNpr1, respectively, with a good fit of the αI-gliadin substrate inside the putative active clefts (Fig. 6A). The atomic contact energy (ACE) of m-NrNpr1 and m-NvNpr were -193.76 and -182.07, respectively. Furthermore, the approximate interface area of the docking complex with m-NvNpr and m-NrNpr1 were 992.60 and 1049.40, respectively. This supports that both m-NvNpr and m-NrNpr1 could bind αI-gliadin for hydrolysis.

**Figure 5.**
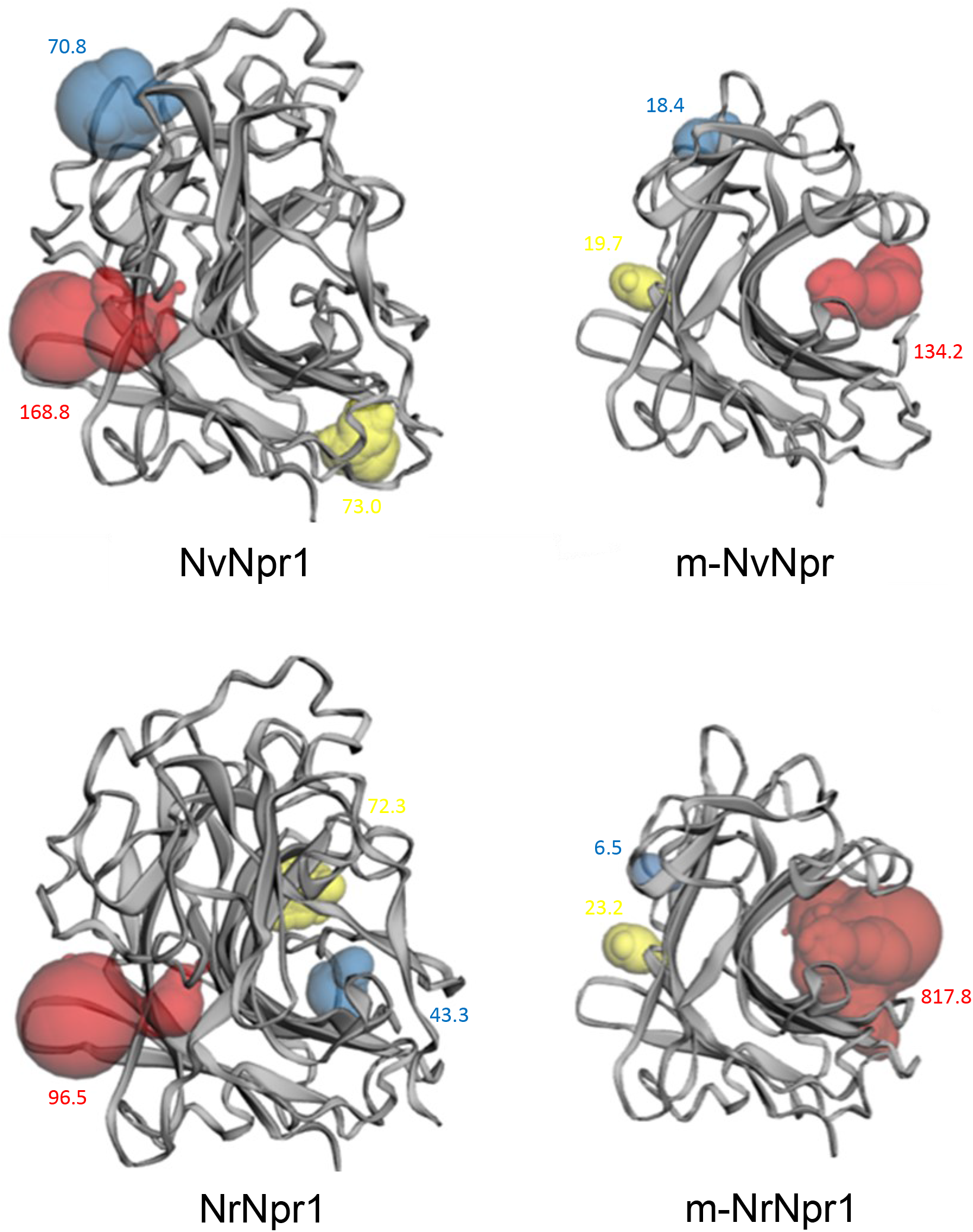
The predicted binding pockets in neprosin proenzymes (NvNpr and NrNpr1) and mature neprosin (m-NvNpr and m-NrNpr1) models. The red pockets have the largest solvent-accessible volume, followed by the yellow and blue pockets. Richard’s solvent-accessible volumes are annotated next to the pockets with the respective colored font.

**Figure 6.**
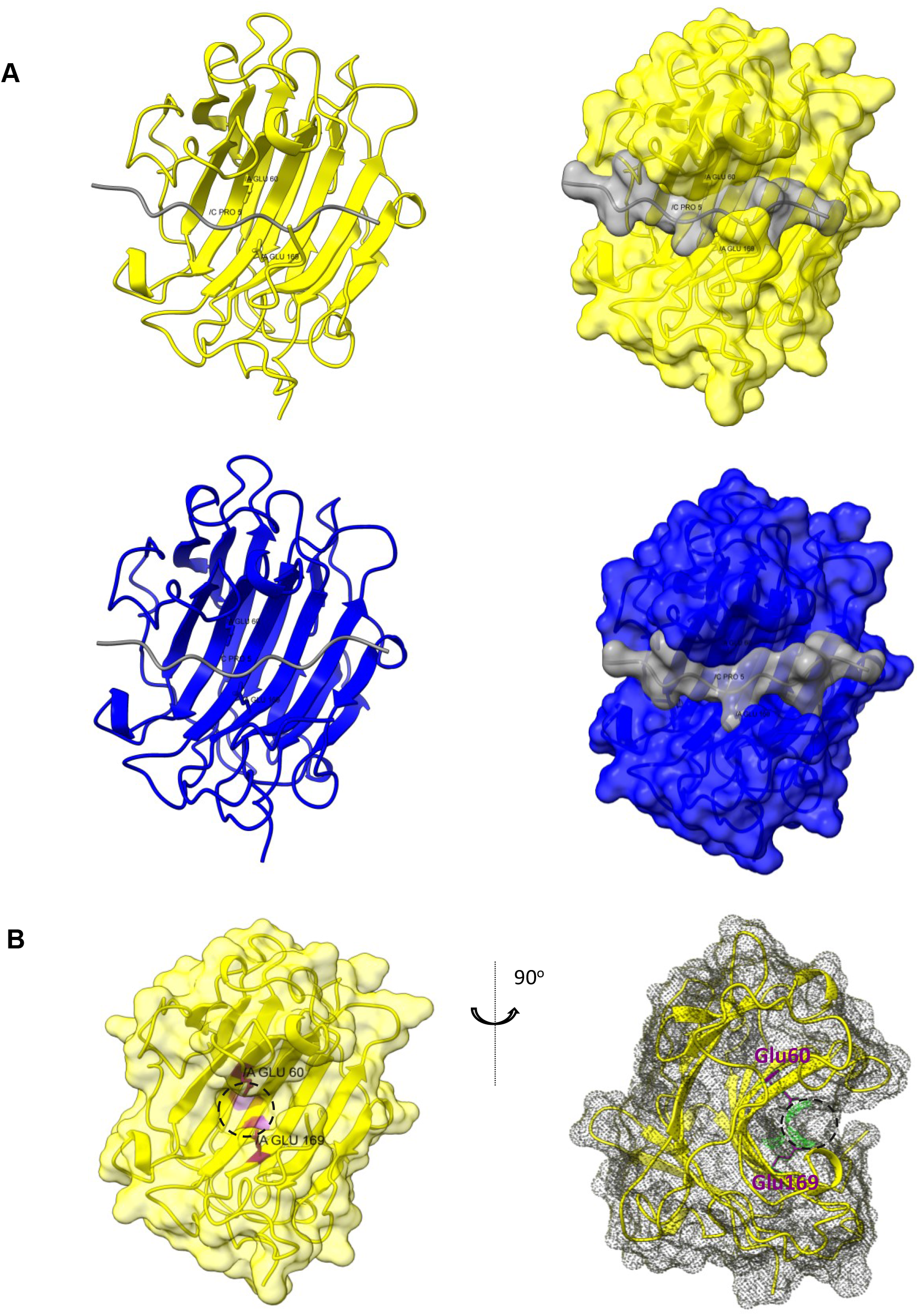
Docking and active cleft analyses of mature neprosins. (A) Top docking solutions of mature neprosins from *N*. × *ventrata* (m-NvNpr) and *N. rafflesiana* (m-NrNpr1) with deaminated αI-gliadin using PatchDock. Left docking models are in ribbon style while models on the right are in a surface view. (B) Surface and ribbon views of m-NvNpr with the putative catalytic dyad (Glu60 and Glu169) in the concavity (dashed circle) of the putative active cleft.

A closer examination revealed Glu60 and Glu169 to be in a concavity located in the putative active cleft (Fig. 6B). The concavity could be the putative active site of the neprosin as it resembles the active site cavities reported in the *Streptomyces sioyaensis* endo-1,3-β*-*glucanase (PDB id: 3dgt) (Hong et al. 2008) and SGP (Pillai et al. 2007). Strikingly, the docking models of m-NvNpr and m-NrNpr1 showed that the helix/turn motif of Pro5 in the αI-gliadin faced towards the putative active cleft and fits into the putative active site cavity containing the putative catalytic dyad (Glu60 and Glu169) of m-NvNpr and m-NrNpr1 (Fig. 6A). The active site cavity may be the substrate recognition S1’ subsite which recognizes proline residues in the substrate, thus providing the preference of proline in P1 for the mature neprosins. The molecular dynamics simulation using CABS-flex 2.0 showed that the putative catalytic dyad (Glu60 and Glu169, numbering in mature neprosin) in both m-NvNpr and m-NrNpr1 were in a stable region with low RMSF (Table 3, Fig. S9).

**Table 3.** RMSF (Å) of the putative catalytic residues in mature neprosin models (m-NvNpr and m-NrNpr1). Residue fluctuation, RMSF(Å) values are extracted from residue fluctuation profile output of molecular dynamics stimulation by CABS-flex 2.0.

| Models | Putative catalytic residue | RMSF (Å) |
| --- | --- | --- |
| m-NvNpr | Glu60 | 0.262 |
|  | Glu169 | 0.373 |
| m-NrNpr1 | Glu60 | 0.181 |
|  | Glu169 | 0.274 |

On the other hand, Glu293 (Glu165 in mature NvNpr) in the binding pocket is not chosen as a candidate catalytic residue for several reasons. Based on the ConSurf analysis (Fig. 2) and the representative proteome 15 (RP15) of the neprosin domain (Fig. S6), Glu293 is not functional and less conserved than Glu188 and Glu296. Secondly, Glu293 is superposed to Val169 of the SMoV peptidase unit, which is not catalytic. In the mature NvNpr, the distance between Glu165 and Glu169 (7.603 Å) is farther than the distance between Glu60 and Glu169 (5.225 Å) (Fig. S10), which is more similar to 4.8 Å between Gln24 and Glu110 catalytic dyad in AGP (Sasaki et al., 2004) and 5.03 Å between Gln53 and Glu136 in SGP. In addition, Glu165 does not form a concavity compared to Glu60 and Glu169 in the putative substrate-binding pocket (Fig. S10). Hence, Glu60 and Glu 169 are more plausible candidates for the catalytic dyad. However, we cannot exclude the possibility that Glu165 could confer catalytic activity since it is in a highly stable position (Fig. S9) and exposed in the putative active cleft of mature NvNpr (Fig. S10). Further experiments will be needed to ascertain the functionality of all three glutamic acid residues.

### 3.6. A proposed model of neprosin regulation and catalytic mechanism

Based on the previous literature and the hypothesis that neprosin belongs to the glutamic peptidase family with autoproteolytic maturation, we proposed a general model for the regulation of NvNpr (Fig. 7). Neprosin and other proteases are expressed in the pitcher tissues even before the pitchers open (Goh et al. 2020). Neprosin proteins undergo post-translational modifications, such as glycosylation and the removal of the signal peptide before being secreted into the pitcher fluid as a proenzyme upon pitcher opening. Their expression and secretion can be induced by ammonium, chitin, and protein from prey in the pitcher fluid (Lee et al. 2016; Wan Zakaria et al. 2019; Zulkapli et al. 2021). The presence of prey (ammonium, chitin, or protein) in the pitcher fluid triggers the acidification of pitcher fluid (Bazile et al. 2015; Lee et al. 2016; Saganová et al. 2018), resulting in the activation of neprosin via autoproteolysis or hydrolysis by other proteases. Autoproteolysis is more likely since the purified recombinant NvNpr can undergo *in vitro* activation (Schräder et al. 2017). Intramolecular autoproteolysis by neprosin itself is possible due to the location of the PP/S cleavage site (Fig. 1) for neprosin maturation is ideally positioned in the flexible loop near the putative active cleft. Furthermore, the N→ O acyl shift could play a role in neprosin maturation by the hydrolysis of the scissile peptide bond of Pro-Ser in the PP/S cleavage site. The nucleophilic serine hydroxyl group could attack the carbonyl group of proline resulting in the formation of an ester. The ester bond is more susceptible to hydrolysis compared to the amide bond but the equilibrium of the N→ O acyl shift is in favor of the amide under physiological conditions (Paulus 2000). However, the acidic condition in the pitcher fluids could shift the equilibrium towards the acidic hydrolysis of ester bonds. Moreover, the tight turn in the loop containing the PP/S cleavage site may cause conformation strain that accelerates the N→ O acyl shift for autoproteolysis, similar to that reported in sea urchin sperm protein, enterokinase, and agrin (SEA) domain autoproteolysis (Sandberg et al. 2008). Hence, the N→ O acyl shift at low pH conditions could be the most likely mechanism for the autoproteolysis of neprosin. During proteolysis, hydrolysis at the disordered region between the Neprosin_AP and neprosin domains in the proenzyme forms the mature neprosin with only the neprosin domain. The mature neprosin structure has an accessible putative active cleft for substrate binding.

**Figure 7.**
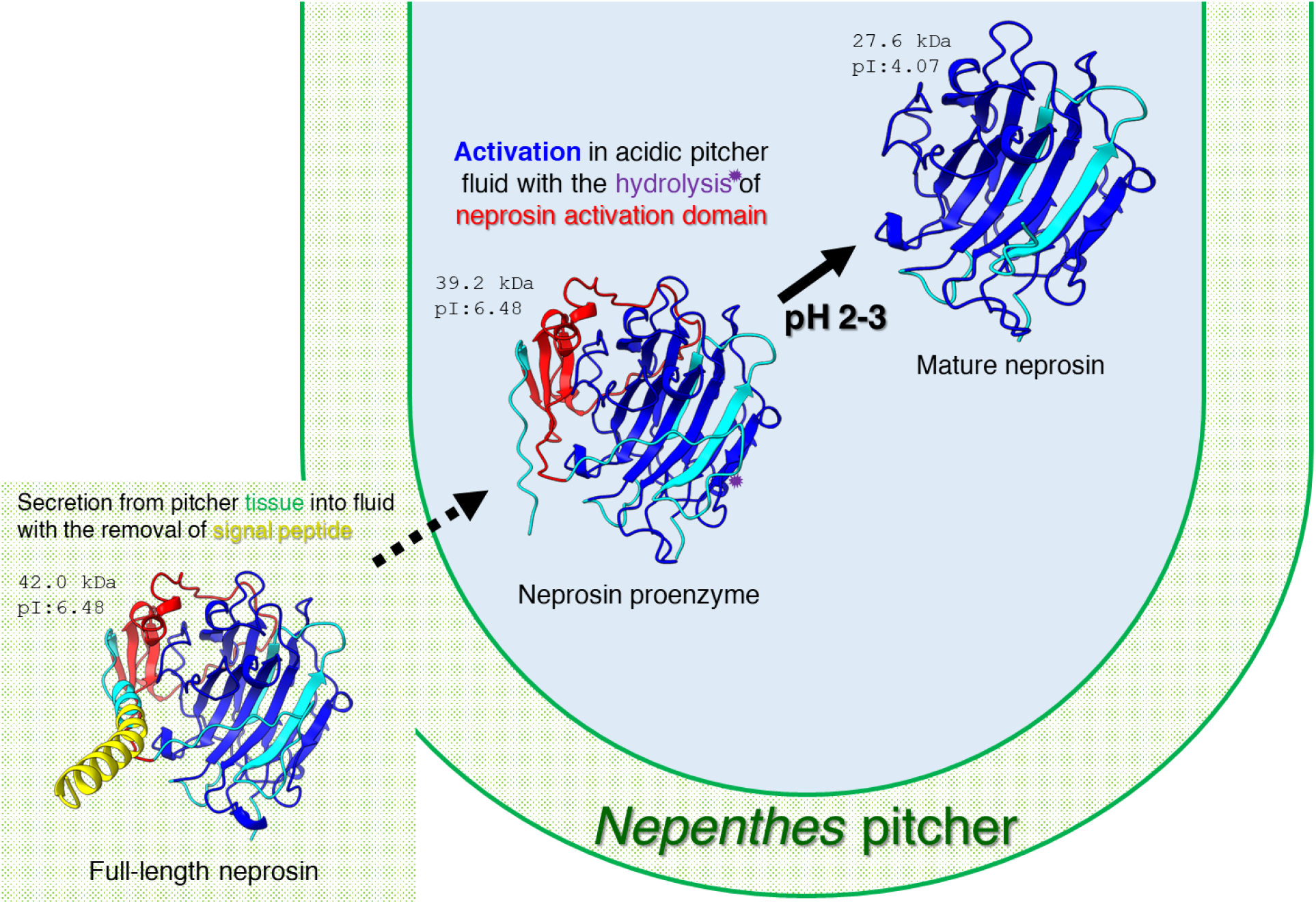
A proposed regulatory mechanism of neprosin protein in the *Nepenthes* pitcher. Theoretical molecular weight (kDa) and isoelectric point (pI) of NvNpr are shown. Mature neprosin is based on the second putative start site as indicated by the purple shape. Different domains of neprosin correspond to different colored fonts.

The proposed catalytic neprosin domain with prolyl endopeptidase activity (Rey et al. 2016) resembles a glucanase-like β-sandwich, which comprises two seven-stranded antiparallel β-sheet, unique to the glutamic peptidase family. The catalytic dyad of two glutamic acids with post-proline cleaving activity strongly supports that neprosin belongs to a new MEROPS glutamic peptidase family. To date, no proteolysis mechanism has been described that involves a catalytic dyad of two glutamic acids as shown by Mann et al. (2019) to be responsible for the post-proline cleaving activity of SMoV peptidase. Here, we propose a possible catalytic mechanism based on the catalytic dyad of AGP shown by Sasaki et al. (2005) with the glutamine (Gln) residue replaced by glutamic acid (Glu) in neprosin (Fig. 8). Since glutamic acid is often found in the protein active binding sites, the hydroxyl group of Glu, despite being negatively charged, could function similarly to that of the Gln amide sidechain in providing electrophilic assistance and oxyanion stabilization to the tetrahedral intermediate state of the peptide bond. Glu169 in mature neprosin could act as a general acid during the first phase of proteolysis by donating a proton to the oxygen of the carbonyl group of the scissile peptide bond. Then, Glu169 acts as a general base to activate a water molecule that carries out a nucleophilic attack on the carbonyl carbon atom of the scissile peptide bond. The water molecule with a hydrogen bond formed with the hydroxyl group of Glu60 donates an OH-group to the carbonyl carbon of the scissile peptide bond to form a tetrahedral intermediate. The process of electron transfer is aided by another water molecule held by the hydrogen bonds with Glu60 and Glu169. The transition state of the scissile peptide bond is stabilized by the hydrogen bonds with the two catalytic Glu residues. In the next phase, the protonated Glu169 donates the proton to the nitrogen of the amide group of the peptide bond. Finally, the protonation of the leaving-group nitrogen triggers the breakdown of the tetrahedral intermediate and causes the peptide bond hydrolysis.

**Figure 8.**
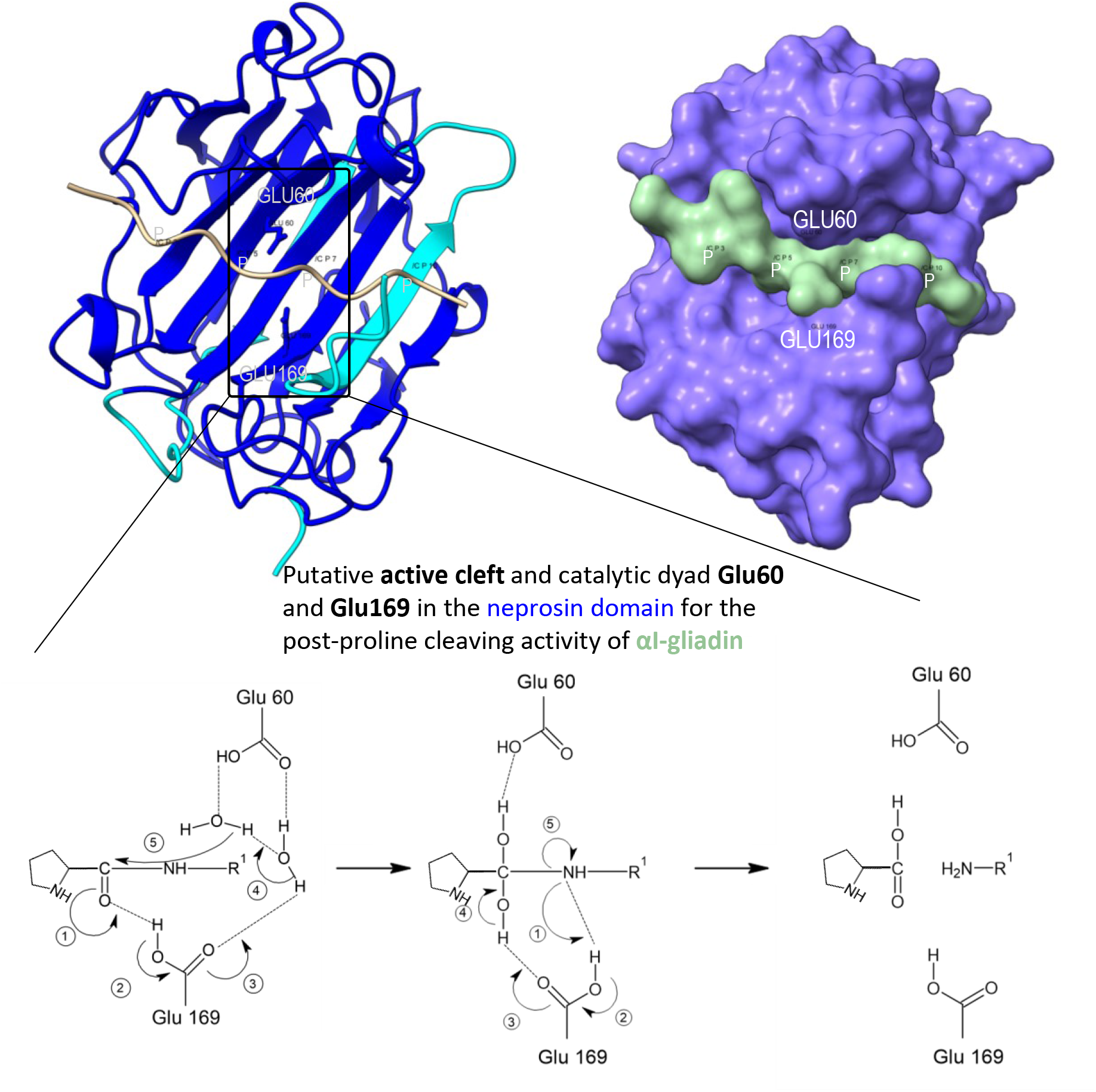
A proposed catalytic mechanism of neprosin inside the active site of substrate binding cleft. Dotted lines indicate hydrogen bonds. Numbers in circle represents sequential reactions.

## 4. Conclusion

In this study, *in silico* structure-function analysis of the neprosin family provided evidence for the classification of neprosins as glutamic peptidases with prolyl endopeptidase activity. It also revealed the putative regulation and catalytic mechanism of neprosin as depicted in Figs. 7 and 8. This is supported by the structural similarity, autoproteolysis activation at low pH conditions, and the presence of glutamic acid residues in the putative active cleft. Moreover, the catalytic dyad of family G3 with post-proline cleaving activity is superposed to the hypothetical catalytic dyad of NvNpr. Further validation experiments such as crystallography studies and the expression of recombinant neprosin with site-directed mutagenesis of the glutamic acid residues will ascertain the functions of these catalytic residues in the neprosin family. It will also be interesting to study the functional divergence of neprosins in other plant species.

## Ethics approval and consent to participate

Not applicable.

## Human and animal rights

No Animals/Humans were used for studies that are the basis of this research.

## Consent for publication

Not applicable.

## Availability of data and materials

All PDB files of protein structures and docking described in this manuscript can be accessed on Figshare (https://doi.org/10.6084/m9.figshare.19187252).

## Acknowledgments

This work was supported by the Malaysian Ministry of Higher Education *Fundamental Research Grant Scheme* (FRGS/1/2019/STG05/UKM/02/10) and Universiti Kebangsaan Malaysia Research University Grant (DIP-2020-005). The group also acknowledges the great effort of the ColabFold team in making the structural prediction via AlphaFold2 and RoseTTAFold accessible. The authors are also grateful to the two anonymous reviewers for their constructive comments to improve this manuscript.

## Author contributions

H-H.G. conceived the study and obtained funding. T.Y.T. and A.B. performed the analysis. T.Y.T. and H-H.G. wrote the manuscript. A.B.R and C.L.N. reviewed the manuscript. All authors approved the final manuscript.

## Supportive/supplementary material

Supplementary material is available on the publisher’s website along with the published article. All PDB files of protein structures and docking described in this manuscript can be accessed on Figshare (https://doi.org/10.6084/m9.figshare.19187252).

## List of Supplementary Table

**Table S1**. Secondary structures in the predicted models of NvNpr and NrNpr1 based on Jmol FirstGlance analysis.

## List of Supplementary Figures

**Figure S1**. Summary workflow of *in silico* analysis on neprosin amino acid sequences.

**Figure S2**. Phylogenetic tree of neprosin amino acid sequences with respective top 10 BLASTP hits from NCBI nr database, constructed using MEGAX package with Maximum Likelihood of 500 bootstraps. The black boxes indicate the neprosin query sequences for BLASTP searches.

**Figure S3**. Phylogenetic tree of neprosins with BLASTP hits against the *Arabidopsis thaliana* genome sequence database using Dicots PLAZA 5.0.

**Figure S4**. Multiple sequence alignment of neprosins and their closely related amino acid sequences of BLASTP hits against NCBI nr (*N. alata* C-terminal peptidases: BAW35438.1 and BAW35437.1) and *Arabidopsis* sequence (AT5G19170 and AT3G48230) using Clustal Omega. The neprosin activation and neprosin domains are highlighted in the red and blue boxes, respectively.

**Figure S5**. The superimposition of scytalidoglutamic peptidase (SGP, 2ifw) and aspergilloglutamic peptidase (AGP, 1y43) crystal structures with the AlphaFold2 neprosin models of *N*. × *ventrata* (NvNpr) and *N. rafflesiana* (NrNpr1).

**Figure S6**. The representative proteome 15 (RP15) and representative proteome 35 (RP35) of the neprosin domain (PF03080). The glutamic acid residues of the putative catalytic dyad were conserved in >0.6 of 427 sequences in RP15 and 2,067 sequences in RP35.

**Figure S7**. The superimposition of AlphaFold2 models of strawberry mottle virus (SMoV) glutamic peptidase unit with neprosins from *N*. × *ventrata* (NvNpr) and *N. rafflesiana* (NrNpr1).

**Figure S8**. The **s**uperimposition and mTM-align pairwise alignment of crystal structures of scytalidoglutamic peptidase (SGP, PDB id: 2ifw) and aspergilloglutamic peptidase (AGP, PDB id: 1y43-B) with the AlphaFold2 models of *N*. × *ventrata* (NvNpr), *N. rafflesiana* (m-NrNpr1), and strawberry mottle virus glutamic peptidase (SMoV, MER1365461). Pairwise TM-score, RMSD, and alignment length (L_ali_) were obtained from the structural alignment using mTM-align, whereas sequence identity (%ID) was based on pairwise sequence alignment using Clustal Omega.

**Figure S9**. The output of molecular dynamics simulation of m-NvNpr (top) and m-NrNpr1 (bottom) by CABS-flex 2.0. (A) three-dimensional visualization of 10 models. (B) Residue fluctuation profile (RMSF) plots.

**Figure S10**. The relative positions and distances between the candidate glutamic acid residues in the putative active cleft. Glu60 and Glu169 can form a subpocket within the putative substrate-binding pocket as outlined in cyan.

