## Supplementary File for "Neprosin belongs to a new family of glutamic peptidase based on *in silico* evidence"

### Supplementary Table

**Table S1**. Secondary structures in the predicted models of NvNpr and NrNpr1 based on Jmol FirstGlance analysis.

### Supplementary Figures

**Figure S1**. The summary workflow of *in silico* analysis on neprosin amino acid sequences.

**Figure S2**. Phylogenetic tree of neprosin amino acid sequences with respective top 10 BLASTP hits from NCBI nr database, constructed using MEGAX package with Maximum Likelihood of 500 bootstraps. The black boxes indicate the neprosin query sequences for BLASTP searches.

**Figure S3**. Phylogenetic tree of neprosins with BLASTP hits against the *Arabidopsis thaliana* genome sequence database using Dicots PLAZA 5.0.

**Figure S4**. Multiple sequence alignment of neprosins and their closely related amino acid sequences of BLASTP hits against NCBI nr (*N. alata* C-terminal peptidases: BAW35438.1 and BAW35437.1) and *Arabidopsis* sequence (AT5G19170 and AT3G48230) using Clustal Omega. The neprosin activation and neprosin domains are highlighted in the red and blue boxes, respectively.

**Figure S5**. The superimposition of the AlphaFold2 neprosin models of *N.* × *ventrata* (NvNpr) and *N. rafflesiana* (NrNpr1), scytalidoglutamic peptidase (SGP, PDB id: 2ifw) and aspergilloglutamic peptidase (AGP, PDB id: 1y43) crystal structures using the 'Matchmaker' function of ChimeraX.

**Figure S6**. The representative proteomes 15 (RP15) and representative proteomes 35 (RP35) of the neprosin domain (PF03080). The glutamic acid residues of the putative catalytic dyad were conserved in >0.6 of 427 sequences in RP15 and 2,067 sequences in RP35.

**Figure S7.** The superimposition of AlphaFold2 models of strawberry mottle virus (SMoV) glutamic peptidase unit with neprosins from *N.* × *ventrata* (NvNpr) and *N. rafflesiana* (NrNpr1).

**Figure S8.** The superimposition and mTM-align pairwise alignment of crystal structures of scytalidoglutamic peptidase (SGP, PDB id: 2ifw) and aspergilloglutamic peptidase (AGP, PDB id: 1y43-B) with the AlphaFold2 models of *N.* × *ventrata* (m-NvNpr), *N. rafflesiana* (m-NrNpr1), and strawberry mottle virus glutamic peptidase (SMoV, MER1365461). Pairwise TM-score, RMSD, and alignment length ($L_{ali}$) were obtained from the structural alignment using mTM-align, whereas sequence identity (%ID) was based on pairwise sequence alignment using Clustal Omega.

**Figure S9.** The output of molecular dynamics simulation of m-NvNpr (top) and m-NrNpr1 (bottom) by CABS-flex 2.0. (A) three-dimensional visualization of 10 models. (B) Residue fluctuation profile (RMSF) plots.

**Figure S10.** The relative positions and distances between the candidate glutamic acid residues in the putative active cleft. Glu60 and Glu169 can form a subpocket within the putative substrate binding pocket as outlined in cyan.

**Table S1**. Secondary structures in the predicted models of NvNpr and NrNpr1 based on Jmol FirstGlance analysis.

| **Jmol attribute** | | **NvNpr** | | **NrNpr1** | |
|---|---|---|---|---|---|
| | | **RoseTTAFold** | **AlphaFold2** | **RoseTTAFold** | **AlphaFold2** |
| **Total amino acids** | | 356 | | | |
| **Total atoms** | | 2,773 | | | |
| **Secondary structure (%)** | | | | | |
| Helices | α | 2.8 | 1.1 | 4.2 | 1.1 |
| | $3_{10}$ | 8.7 | 7.6 | 6.7 | 7.6 |
| | pi | 0 | 0 | 0 | 0 |
| | Total | 11.5 | 8.7 | 10.9 | 8.7 |
| β-strands | | 34.3 | 34.3 | 31.7 | 34.3 |
| Turns | | 16.3 | 13.5 | 14.6 | 12.9 |
| Coil | | 37.9 | 43.5 | 42.7 | 44.1 |
| Jmol visualization | | 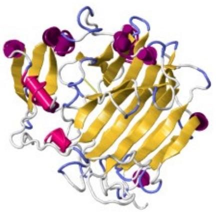 | 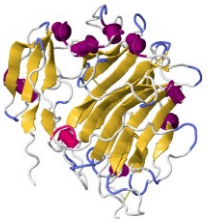 | 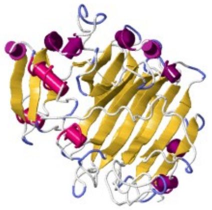 | 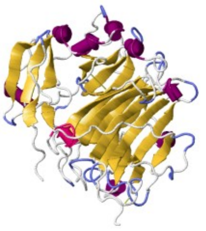 |

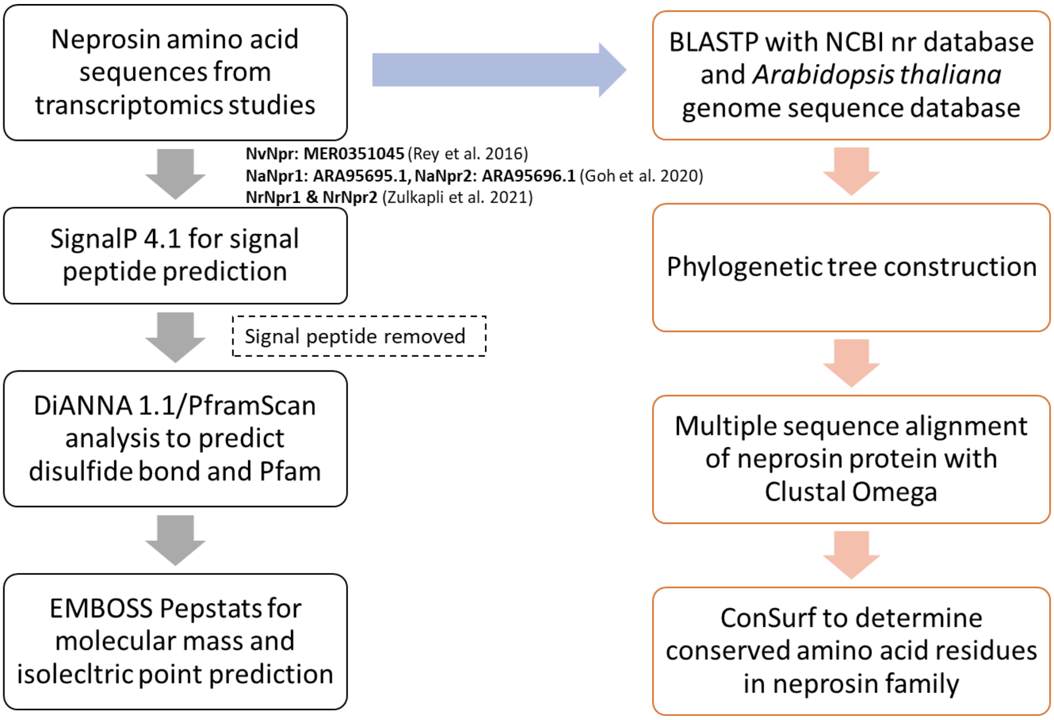


**Figure S1**. The summary workflow of *in silico* analysis on neprosin amino acid sequences.

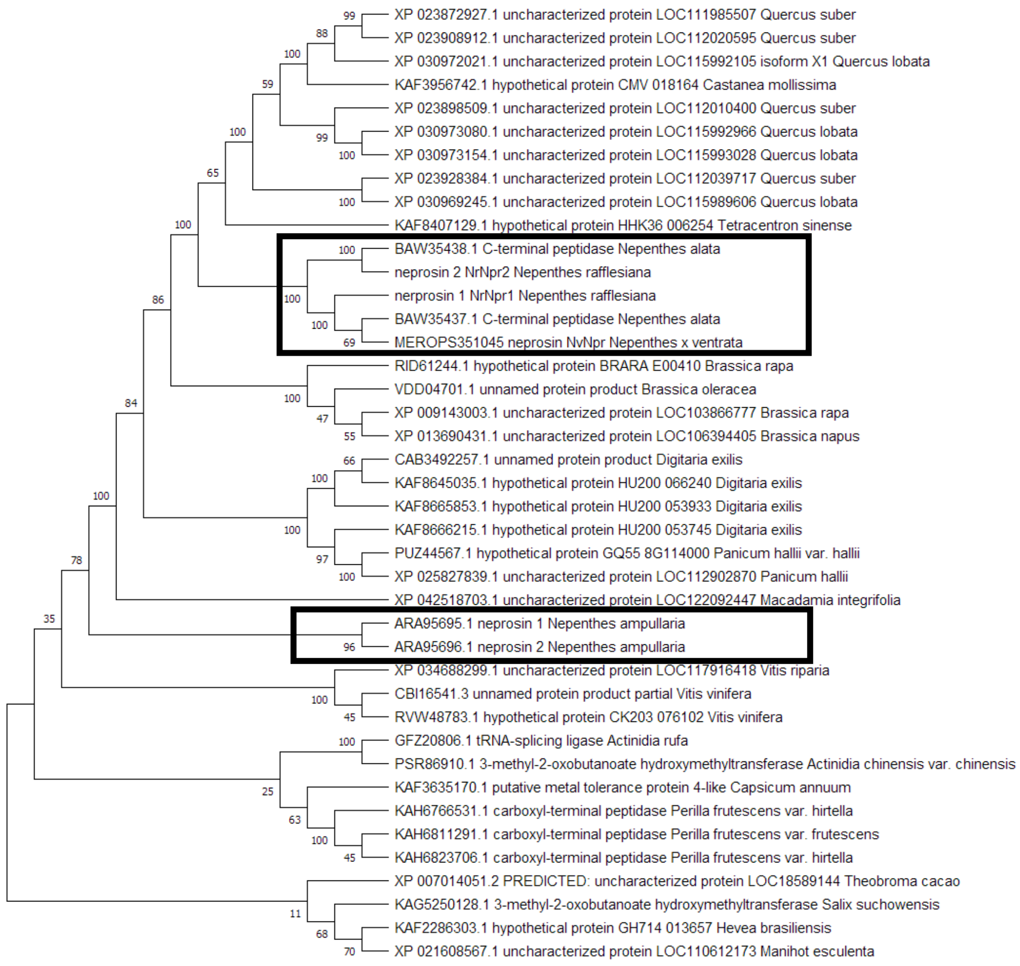


**Figure S2**. Phylogenetic tree of neprosin amino acid sequences with respective top 10 BLASTP hits from NCBI nr database, constructed using MEGAX package with Maximum Likelihood of 500 bootstraps. The black boxes indicate the neprosin query sequences for BLASTP searches.

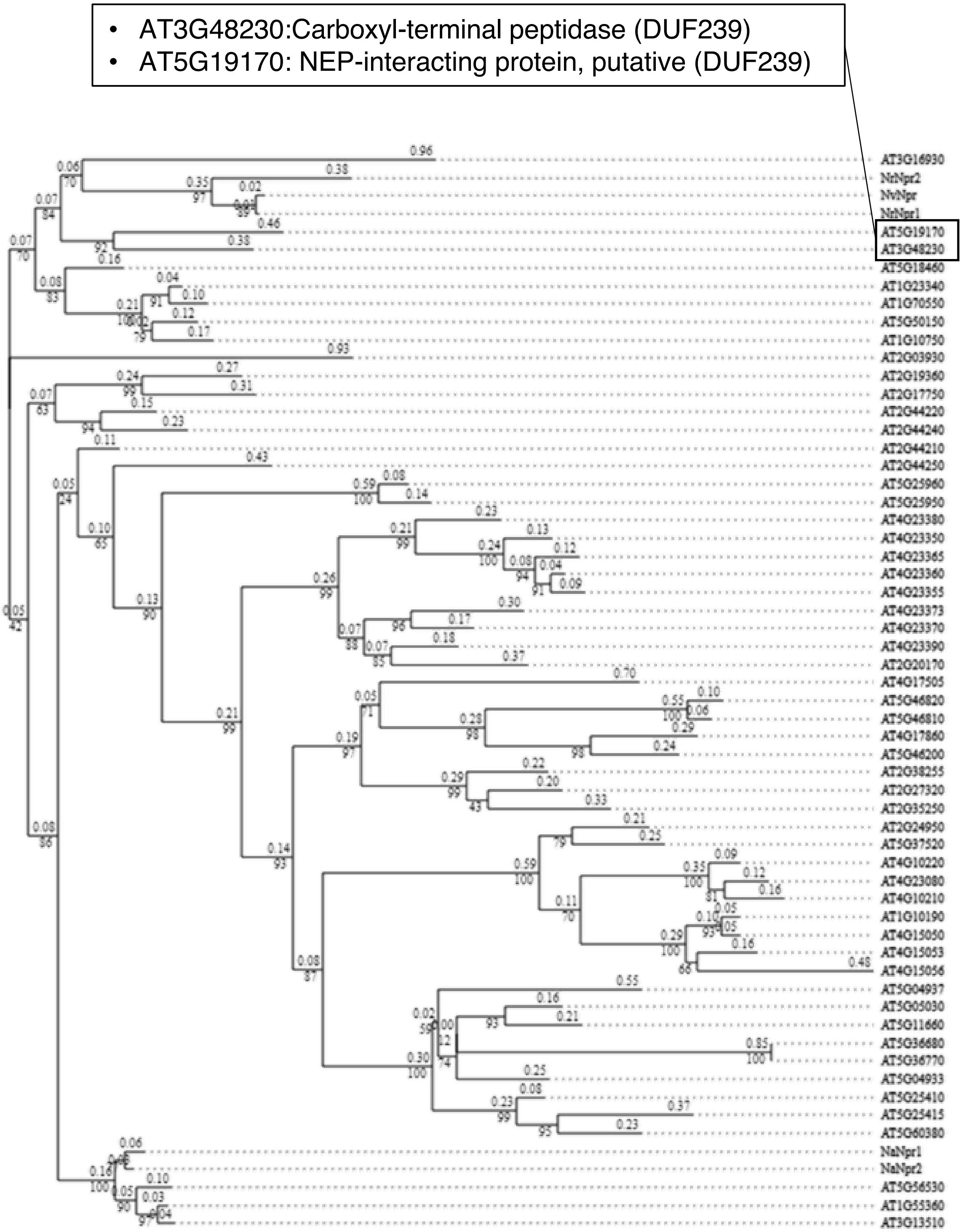


**Figure S3**. Phylogenetic tree of neprosins with BLASTP hits against the *Arabidopsis thaliana* genome sequence database using Dicots PLAZA 5.0.

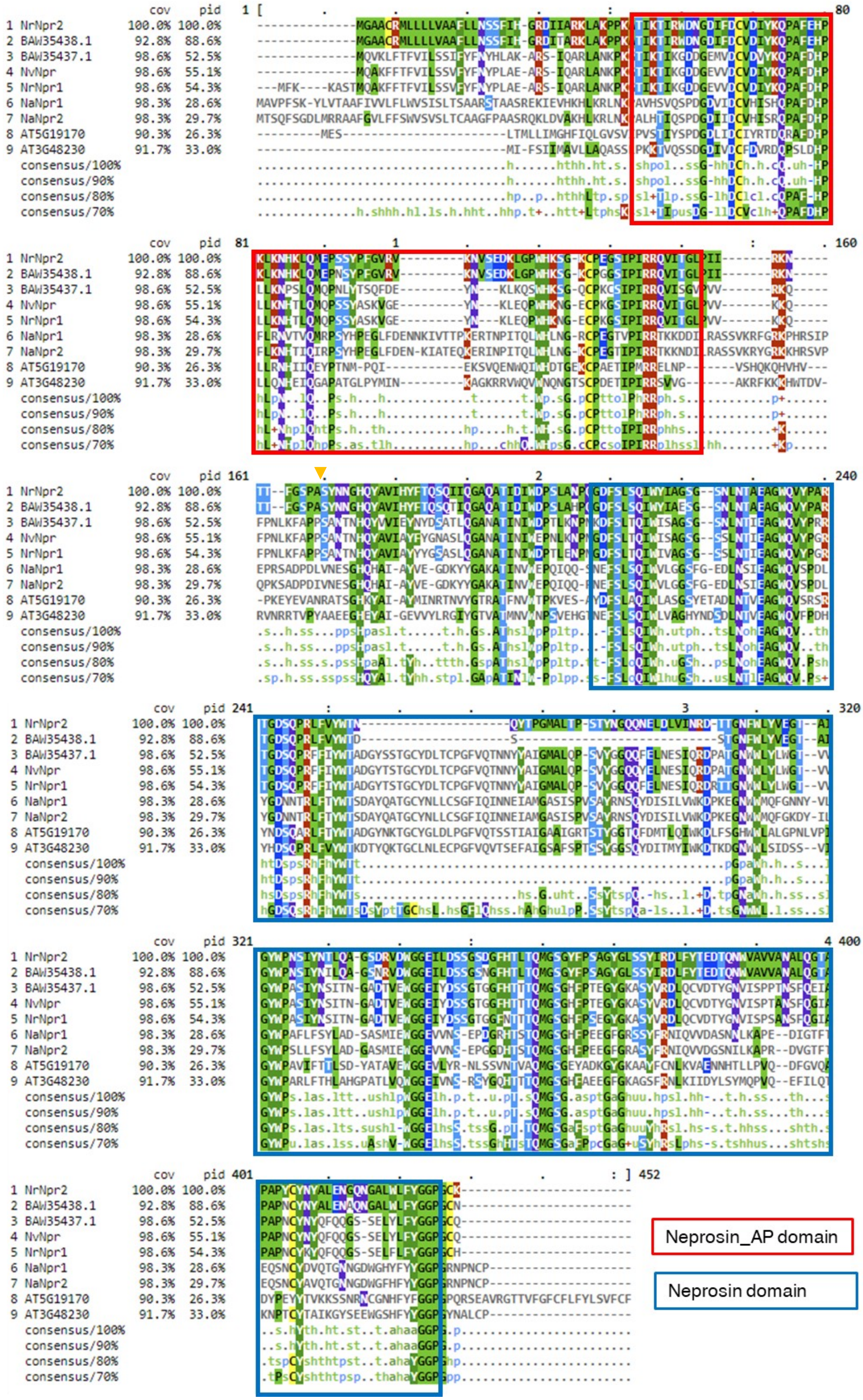


**Figure S4**. Multiple sequence alignment of neprosins and their closely related amino acid sequences of BLASTP hits against NCBI nr (*N. alata* C-terminal peptidases: BAW35438.1 and BAW35437.1) and *Arabidopsis* sequence (AT5G19170 and AT3G48230) using Clustal Omega. The neprosin activation and neprosin domains are highlighted in the red and blue boxes, respectively. ▼ indicates the proposed second start site of mature NvNpr by Rey et al. (2016).

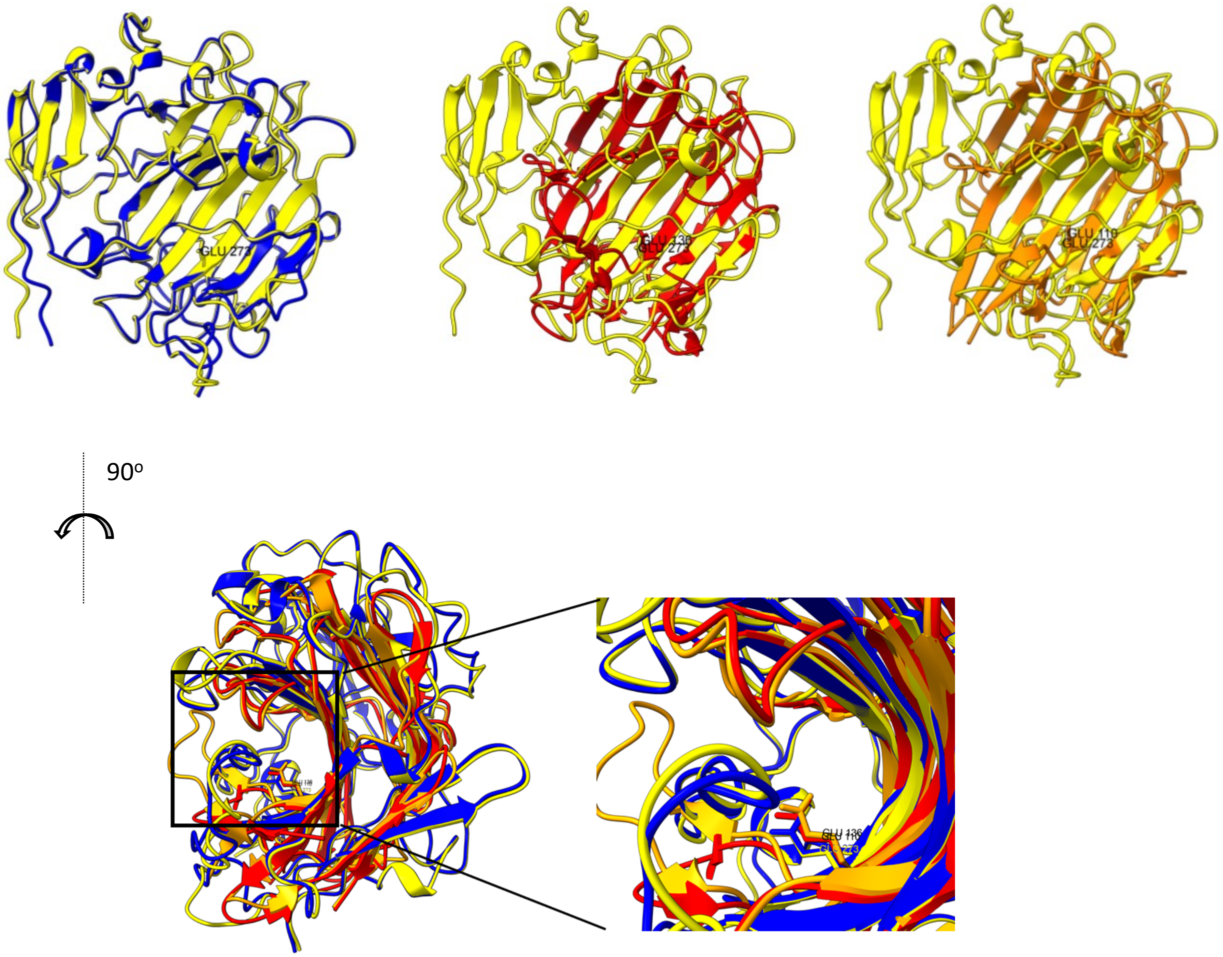


| Model | Superposed Glu position | Note |
|---|---|---|
| 2ifw | 136 | Nucleophile/general acid |
| 1y43 | 110 | Nucleophile/general acid |
| NvNpr | 273 | |
| NrNpr1 | 273 | |

**Figure S5**. The superimposition of the AlphaFold2 neprosin models of *N.* × *ventrata* (NvNpr) and *N. rafflesiana* (NrNpr1), scytalidoglutamic peptidase (SGP, PDB id: 2ifw) and aspergilloglutamic peptidase (AGP, PDB id: 1y43) crystal structures using the 'Matchmaker' function of ChimeraX.

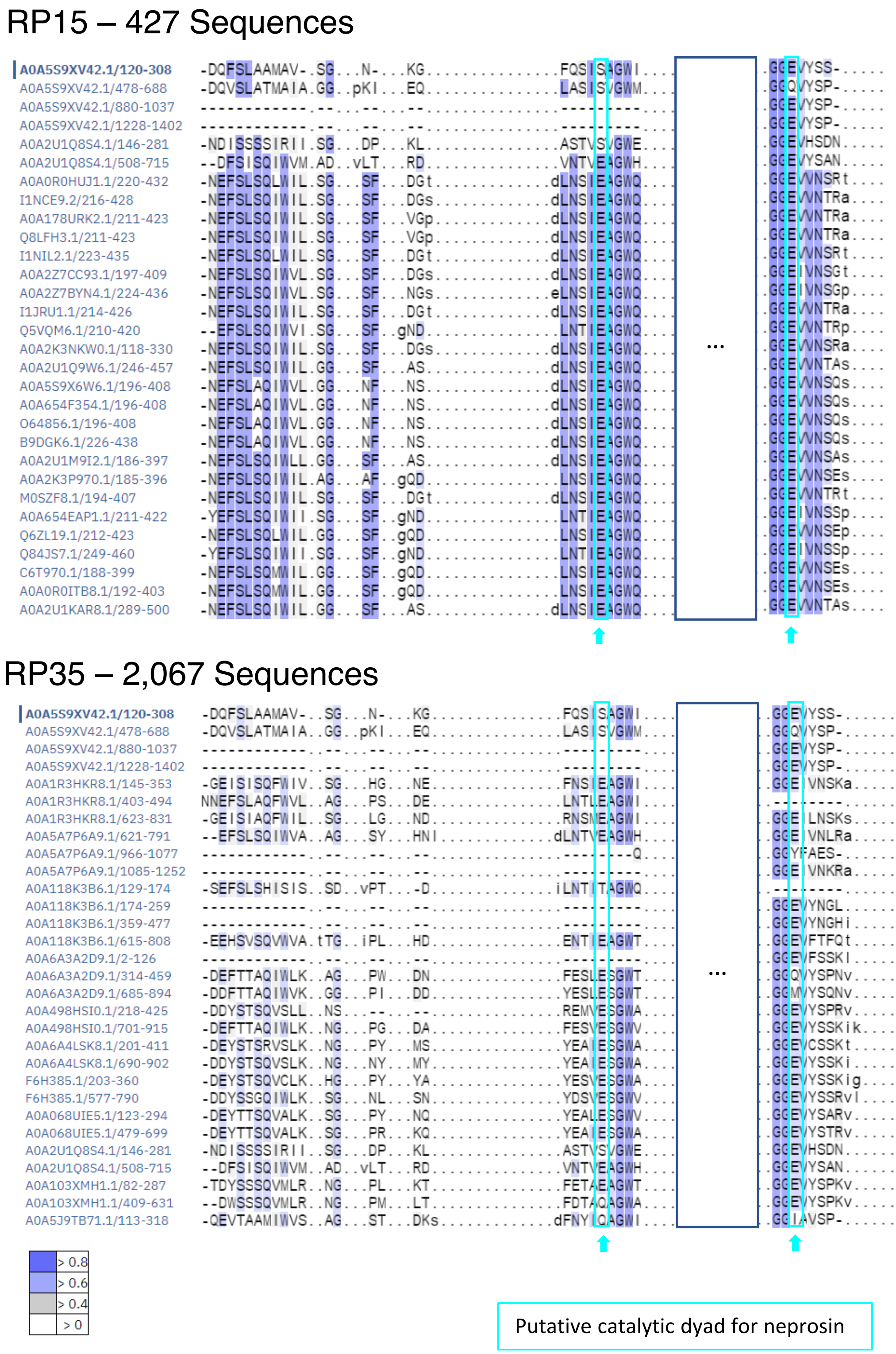


**Figure S6**. The representative proteome 15 (RP15) and representative proteome 35 (RP35) of neprosin domain (PF03080) in the Pfam database. The glutamic acid residues of the putative catalytic dyad were conserved in >0.6 of 427 sequences in RP15 and 2,067 sequences in RP35.

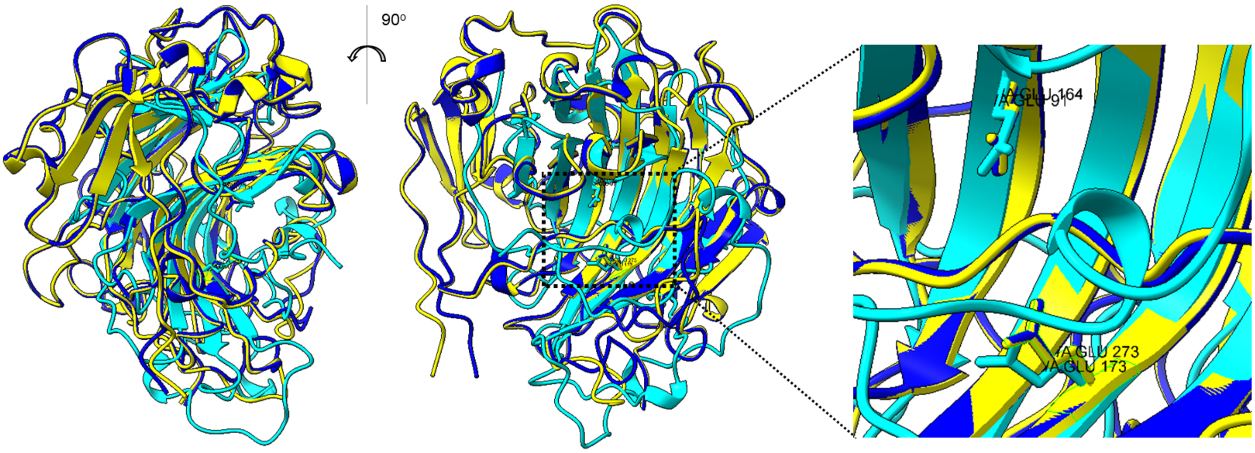


| Model | Superposed GLU position | Note |
|---|---|---|
| SMoV | 91, 173 | Required for protease activity |
| NvNpr | 164, 273 | |
| NrNpr1 | 164, 273 | |

**Figure S7.** The superimposition of AlphaFold2 models of strawberry mottle virus (SMoV) glutamic peptidase unit with neprosins from *N.* × *ventrata* (NvNpr) and *N. rafflesiana* (NrNpr1).

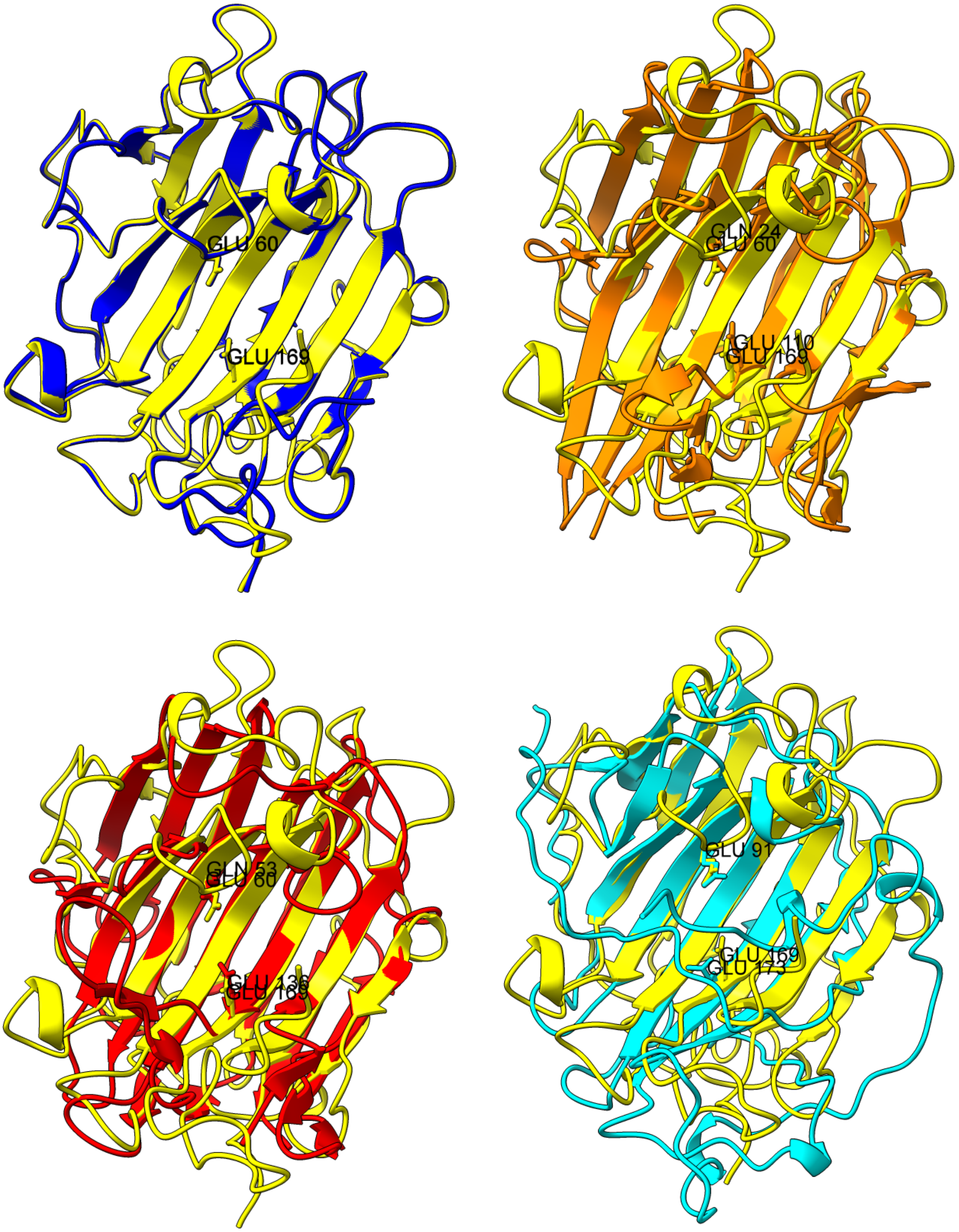


| RMSD / TM-score | AGP-chain B | SGP | m-NvNpr | m-NrNpr1 | SMoV |
|---|---|---|---|---|---|
| AGP-chain B | | 1.230 | 3.230 | 3.392 | 3.618 |
| SGP | 0.9356 | | 3.322 | 3.304 | 3.894 |
| m-NvNpr | 0.7264 | 0.7451 | | 0.908 | 3.258 |
| m-NrNpr1 | 0.7287 | 0.7459 | 0.9867 | | 3.463 |
| SMoV | 0.5508 | 0.5742 | 0.6121 | 0.6069 | |

| %ID / $L_{ali}$ | AGP-chain B | SGP | m-NvNpr | m-NrNpr1 | SMoV |
|---|---|---|---|---|---|
| AGP-chain B | | 44.5 | 8.8 | 8.1 | 14.8 |
| SGP | 166 | | 10.9 | 10.6 | 11.0 |
| m-NvNpr | 128 | 154 | | 94.0 | 11.0 |
| m-NrNpr1 | 128 | 153 | 247 | | 12.2 |
| SMoV | 95 | 114 | 142 | 140 | |

**Figure S8.** The superimposition and mTM-align pairwise alignment of crystal structures of scytalidoglutamic peptidase (SGP, PDB id: 2ifw) and aspergilloglutamic peptidase (AGP, PDB id: 1y43-B) with the AlphaFold2 models of *N.* × *ventrata* (m-NvNpr), *N. rafflesiana* (m-NrNpr1), and strawberry mottle virus glutamic peptidase (SMoV, MER1365461). Pairwise TM-score, RMSD, and alignment length ($L_{ali}$) were obtained from the structural alignment using mTM-align, whereas sequence identity (%ID) was based on pairwise sequence alignment using Clustal Omega.

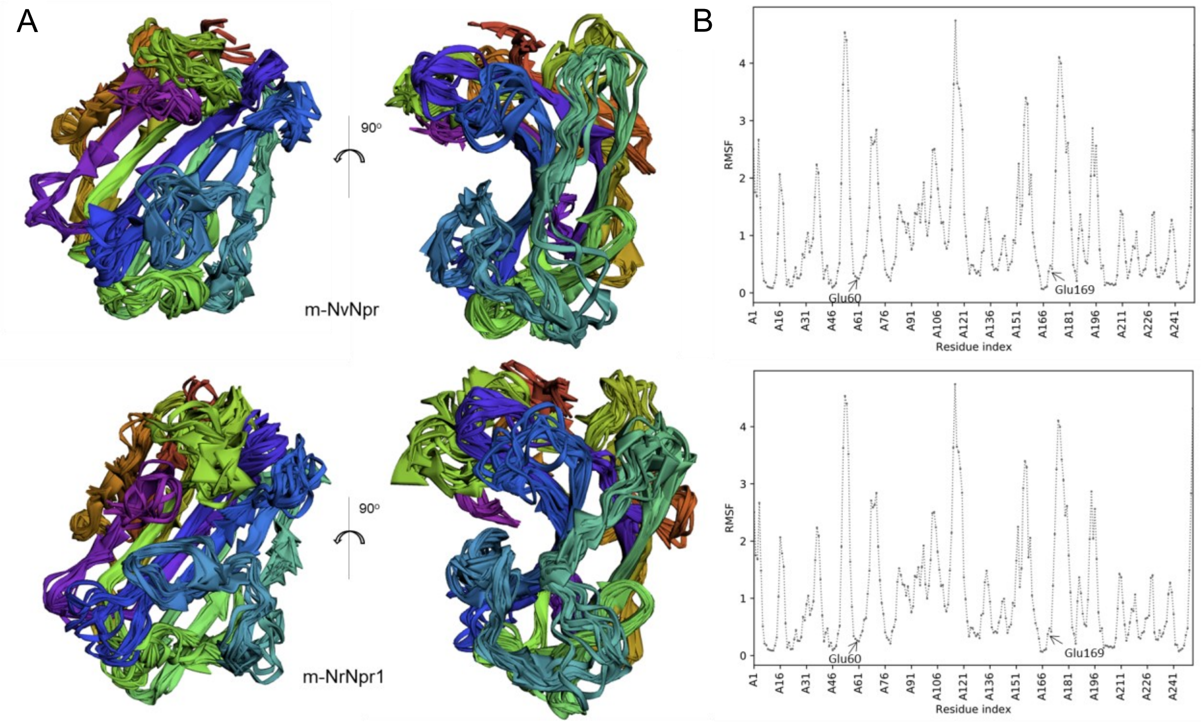


**Figure S9.** The output of the molecular dynamics simulation of m-NvNpr (top) and m-NrNpr1 (bottom) by CABS-flex 2.0. (A) three-dimensional visualization of 10 models. (B) Residue fluctuation profile (RMSF) plots.

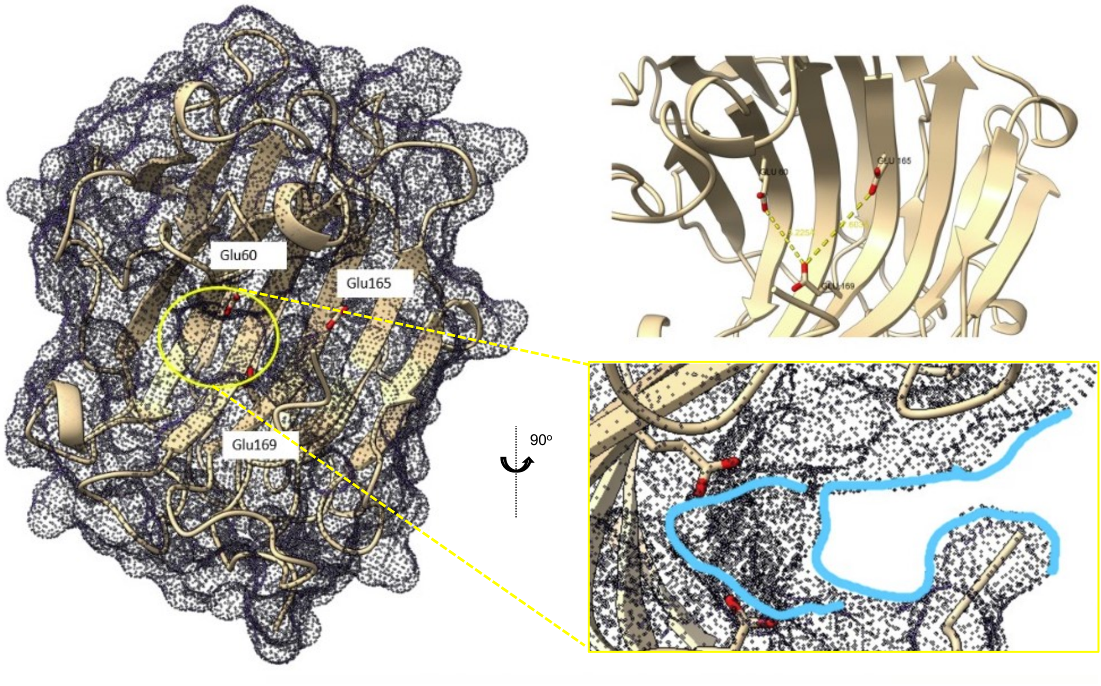


**Figure S10.** The relative positions and distances between the candidate glutamic acid residues in the putative active cleft. Glu60 and Glu169 can form a subpocket within the putative substrate binding pocket as outlined in cyan.
